# Lactylation fuels nucleotide biosynthesis and facilitates deuterium metabolic imaging of tumor proliferation in H3K27M-mutant gliomas

**DOI:** 10.1101/2025.01.02.631150

**Authors:** Georgios Batsios, Céline Taglang, Suresh Udutha, Anne Marie Gillespie, Simon P Robinson, Timothy Phoenix, Sabine Mueller, Sriram Venneti, Carl Koschmann, Pavithra Viswanath

## Abstract

Oncogenes hyperactive lactate production, but the mechanisms by which lactate facilitates tumor growth are unclear. Here, we demonstrate that lactate is essential for nucleotide biosynthesis in pediatric diffuse midline gliomas (DMGs). The oncogenic histone H3K27M mutation upregulates phosphoglycerate kinase 1 (PGK1) and drives lactate production from [U-^13^C]-glucose in DMGs. Lactate activates the nucleoside diphosphate kinase NME1 via lactylation and promotes the synthesis of nucleoside triphosphates essential for tumor proliferation. Importantly, we show that this mechanistic link between glycolysis and nucleotide biosynthesis provides a unique opportunity for deuterium metabolic imaging of DMGs. Spatially mapping ^2^H-lactate production from [6,6-^2^H]-glucose allows visualization of the metabolically active tumor lesion and provides an early readout of response to standard-of-care radiation and targeted therapy that precedes extended survival and reflects pharmacodynamic alterations at the tissue level in preclinical DMG models *in vivo* at clinical field strength (3T). In essence, we have identified an H3K27M-lactate-NME1 axis that promotes DMG proliferation and facilitates non-invasive metabolic imaging of DMGs.

**STATEMENT OF SIGNIFICANCE:** This study establishes a role for lactate in driving nucleotide biosynthesis in DMGs. Importantly, imaging lactate production from glucose using DMI provides a readout of tumor proliferation and early response to therapy in clinically relevant DMG models. Our studies lay the foundation for precision metabolic imaging of DMG patients.

## INTRODUCTION

DMGs are the most lethal form of malignant primary brain cancer in children, with a median overall survival of ∼11 months from initial diagnosis (1,2). These tumors, which include those previously referred to as diffuse intrinsic pontine glioma, arise in delicate anatomical regions such as the pons and thalamus that prevent surgical resection (1,2). Mutation of lysine 27 of histone H3.3 (*H3F3A*) or H3.1 (*HIST1H3B*), most commonly to methionine (H3K27M), is the cardinal oncogenic event in DMGs (1–3). The H3K27M mutation is clonal, retained at recurrence, and functionally relevant since it leads to epigenetic alterations that drive oncogenic gene expression (1–3). The H3K27M mutation disrupts the catalytic activity of the polycomb repressive complex 2 and leads to the global loss of H3K27 trimethylation (H3K27me3), which is associated with the transcriptional repression of gene expression in tumor cells (3). H3K27M also leads to a global increase in H3K27 acetylation (H3K27Ac), an epigenetic mark that is associated with transcriptional activation (3). As a result of these epigenetic alterations, the H3K27M mutation facilitates gene expression that cooperates with mutations in tumor suppressor or receptor tyrosine kinase signaling pathways to initiate tumorigenesis (1–3). Studies show that silencing the H3K27M mutation downregulates the expression of genes associated with stemness and cell cycle progression, decreases proliferation, and delays tumor growth *in vivo* (4,5).

The critical anatomical location of DMGs prevents gross total resection (1,2). Fractionated radiation is the standard of care for DMG patients (1,2). Recently, the imipridones ONC201 and ONC206 have emerged as potential therapies for DMG patients (6,7). Originally identified as antagonists of the dopamine receptor 2 (DRD2), imipridones also function as allosteric agonists of the mitochondrial protease ClpP (6–10). ClpP agonism by imipridones leads to the degradation of key mitochondrial proteins such as the electron transport chain components complex I, II, and IV, and the induction of apoptotic cell death (6–11). Both ONC201 and ONC206 are brain penetrant and significantly extend survival in preclinical DMG models (10,11). Importantly, ONC201 is well tolerated in patients and has shown clinical benefit in a trial in pediatric DMG patients with non-recurrent tumors (median overall survival of 21.7 months vs. 12 months for historical controls) (9). Studies in patients with recurrent tumors also reported potential benefits for ONC201, with a median overall survival of 13.7 months (12).

Magnetic Resonance Imaging (MRI) is the mainstay for assessing disease progression and treatment response in DMG patients (13). Although DMGs, especially those located in the pons, have a characteristic radiographic appearance, they are diffusely infiltrative tumors with indistinct borders (13,14). As such, it is difficult, even for experienced radiologists, to accurately quantify tumor volume by MRI (13,14). Importantly, it is difficult to determine what constitutes a biologically meaningful reduction in tumor volume following the initiation of radiation or experimental therapies such as imipridones (13). As a result, according to current guidelines, multiple MRI scans separated by ≥8 weeks are required to determine if the patient is responding to therapy (13). This lack of reliable response biomarkers hinders the proper stratification of responders from non-responders and leads to devastating consequences such as ineffective treatment for patients with progressive disease and, conversely, unnecessary treatment, and side effects for patients with responsive disease (13). It is also an important clinical research and drug development problem since it hinders the proper interpretation of the efficacy of novel drugs, such as imipridones, in clinical trials. There is an unmet need for novel, clinically translatable, MRI-compatible biomarkers that report on the H3K27M mutation and response to therapy in DMGs.

Oncogenic events rewire glucose metabolism to fuel uncontrolled tumor proliferation (15). Elevated glucose uptake and conversion via glycolysis to lactate generates biosynthetic precursors and ATP and maintains the NAD+/NADH ratio (15,16) (see Fig. 1A). Pyruvate oxidization via the tricarboxylic acid (TCA) cycle produces ATP, NADH, and anaplerotic substrates such as α-ketoglutarate, glutamate, citrate, aspartate, and asparagine, which contribute to lipid and protein synthesis (15,16). Glucose metabolism via the pentose phosphate pathway (PPP) is the major source of NADPH and precursors for nucleotide biosynthesis. Specifically, glucose-6-phosphate is converted to 6-phosphogluconate and ribose-5-phosphate, which is the building block for both purine (AMP, GMP) and pyrimidine (UMP, CTP) nucleotides. Beyond biosynthesis and bioenergetics, glucose-derived metabolites such as acetyl CoA, α-ketoglutarate, and lactate induce post-translational modifications that regulate cell signaling and gene expression (17). For instance, the availability of α-ketoglutarate and acetyl CoA governs histone methylation and acetylation and, thereby, gene expression (17,18). Emerging evidence indicates that lactate-driven lactylation of proteins and histones is a novel post-translational modification that regulates gene expression (19,20). However, whether the H3K27M mutation rewires glycolytic lactate production, and whether lactate and/or lactylation influence tumor proliferation remain unknown.

**Figure 1.**
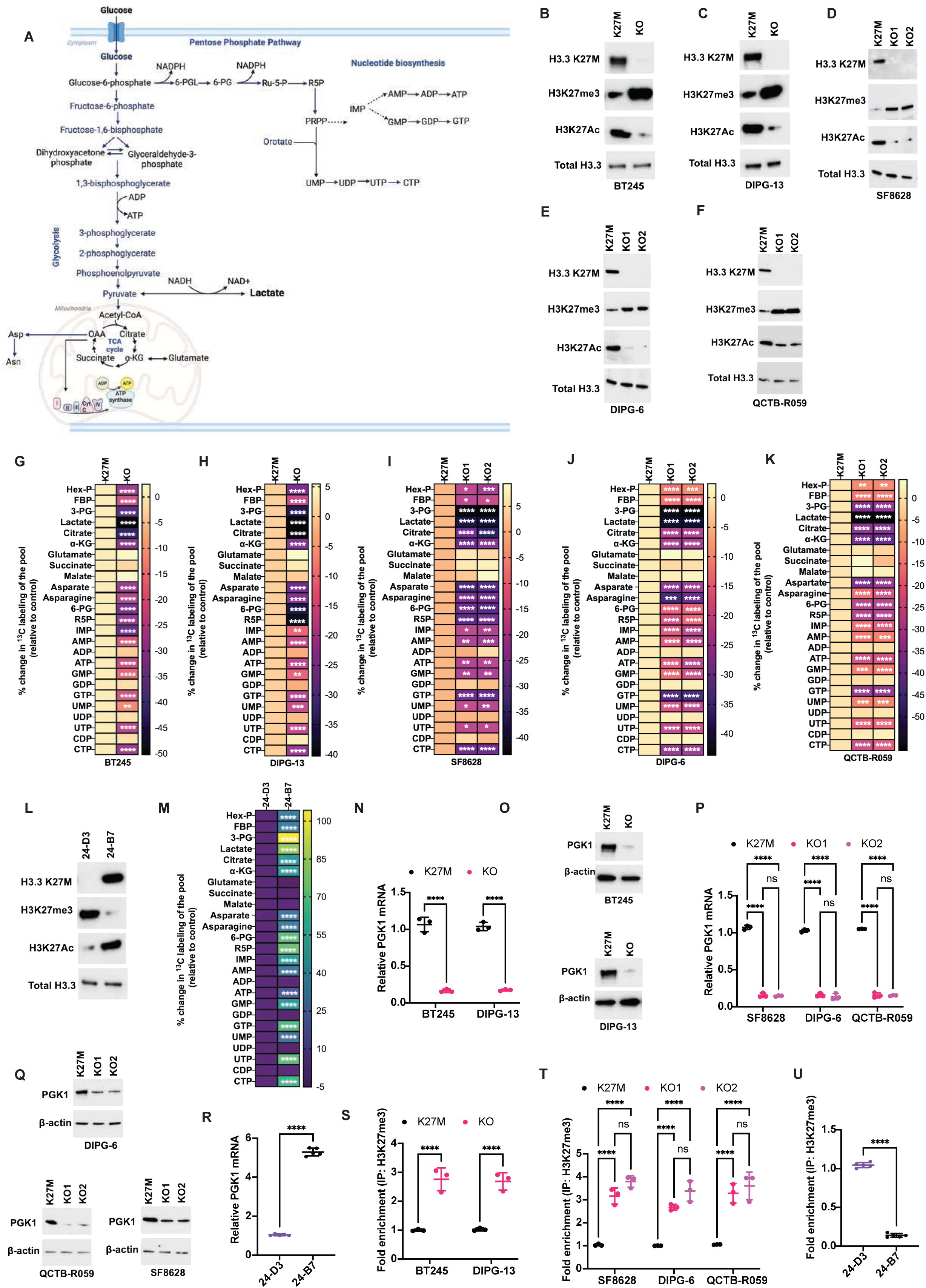
The H3K27M mutation drives glycolytic lactate production in DMGs. **(A)** Schematic illustration of the pathways of glucose metabolism via glycolysis, the TCA cycle, the PPP, and nucleotide biosynthesis. The figure was generated using Biorender. Western blots for mutant H3.3K27M, H3K27me3, H3K27Ac, and total H3.3 in K27M and KO cells for the BT245 **(B)**, DIPG-13 **(C)**, SF8628 **(D)**, DIPG-6 **(E)**, and QCTB-R059 **(F)** models. Quantification of the % change in ^13^C labeling of the metabolite pool from [U-^13^C]-glucose in K27M and KO cells for the BT245 **(G)**, DIPG-13 **(H)**, SF8628 **(I)**, DIPG-6 **(J)**, and QCTB-R059 **(K)** models. Hex-P is hexose monophosphate; FBP is fructose-1,6-bisphosphate; 3-PG is 3-phosphoglycerate; α-KG is α-ketoglutarate; 6-PG is 6-phosphogluconate; R5P is ribose-5-phosphate. **(L)** Western blots for mutant H3.3K27M, H3K27me3, H3K27Ac, and total H3.3 in isogenic H3.3K27M (24-B7) and H3 wild-type (24-D3) cells. **(M)** Quantification of the % change in ^13^C labeling of the metabolite pool from [U-^13^C]-glucose in isogenic H3.3K27M (24-B7) and H3 wild-type (24-D3) cells. PGK1 mRNA **(N)** and protein **(O)** expression in K27M and KO cells for the BT245 and DIPG-13 models. PGK1 mRNA **(P)** and protein **(Q)** expression in K27M and KO cells for the SF8628, DIPG-6, and QCTB-R059 models. **(R)** PGK1 mRNA in 24-B7 and 24-D3 cells. **(S)** Quantification of H3K27me3 enrichment at the PGK1 promoter in K27M and KO cells for the BT245 and DIPG-13 models. **(T)** Quantification of H3K27me3 enrichment at the PGK1 promoter in K27M and KO cells for the SF8628, DIPG-6, and QCTB-R059 models. **(U)** Quantification of H3K27me3 enrichment at the PGK1 promoter in 24-B7 and 24-D3 cells.

The relationship between oncogenic events, metabolic reprogramming, and tumor growth provides a unique opportunity to devise *in vivo* metabolic imaging modalities that signal the presence of an oncogene (21). Magnetic resonance spectroscopy (MRS) is a non-invasive method of imaging tissue metabolism *in vivo* (22). ^1^H-MRS interrogates the nuclear magnetic resonance of protons and provides a readout of steady-state metabolite concentrations (22). However, steady-state metabolite pool sizes do not necessarily reflect metabolic pathway activity. ^13^C-MRS following administration of a ^13^C-labeled substrate such as glucose is the gold standard for quantifying dynamic metabolic activity but is limited by inherently low sensitivity (22). Deuterium metabolic imaging (DMI), which probes the magnetic resonance of the ^2^H nucleus, recently emerged as a novel, clinical-stage method of tracing the metabolism of ^2^H-labeled substrates *in vivo* (23–26). Following administration of [6,6-^2^H]-glucose, glycolytic conversion to lactate vs. oxidation to glutamate can be non-invasively visualized in the brain *in vivo* (23–25). Importantly, [6,6-^2^H]-glucose can be orally administered and has been used to visualize tumor burden in adult glioblastoma patients (23,27).

The goal of this study was to mechanistically dissect the association between the H3K27M mutation, glucose metabolism, and tumor proliferation and to assess if this information can be used for non-invasive imaging of DMGs. Our studies indicate that the H3K27M mutation upregulates the rate-limiting glycolytic enzyme PGK1 and drives [U-^13^C]-glucose metabolism via glycolysis to lactate. Lactate activates the nucleoside diphosphate kinase NME1 via lactylation at lysine 49 (K49) and drives the synthesis of nucleoside triphosphates (NTPs) from nucleoside diphosphates (NDPs), which is essential for tumor proliferation. Importantly, we demonstrate that tracing lactate production from [6,6-^2^H]-glucose provides a non-invasive readout of the H3K27M mutation and facilitates early assessment of tumor burden and treatment response in clinically relevant, patient-derived DMG models *in vivo*.

## RESULTS

### The H3K27M mutation is essential for glucose metabolism via glycolysis the TCA cycle, the PPP, and nucleotide biosynthesis in patient-derived DMG cells

To determine whether the H3K27M mutation reprograms glucose metabolism in a clinically relevant setting, we quantified the effect of the loss of the H3.3 K27M mutation in a diverse panel of patient-derived DMG models (BT245, SF8628, DIPG-6, DIPG-13, and QCTB-R059). For the BT245 and DIPG-13 models, we examined cells in which the H3.3K27M mutation was deleted by CRISPR-Cas9 editing by the Jabado laboratory (28). For the SF8628, DIPG-6, and QCTB-R059 models, we used an orthogonal approach and designed antisense oligonucleotides (ASOs) to silence the H3.3K27M mutation as described earlier (5). In all cases, cells with H3.3K27M expression are referred to as K27M, and cells without H3.3K27M expression are denoted as KO. In line with previous studies (4,5,28), we confirmed that both CRISPR-Cas9-based editing and ASO-based silencing of mutant H3.3K27M expression increased global H3K27me3 and reduced H3K27Ac in all DMG models (Fig. 1B-1F).

We then used stable isotope tracing combined with liquid chromatography-mass spectrometry (LC-MS) to trace [U-^13^C]-glucose metabolism in K27M and KO cells (29). As shown in Fig. 1G-1K, ^13^C labeling of metabolites of glycolysis (hexose phosphate, fructose-1,6-bisphosphate, 3-phosphoglycerate, lactate), the TCA cycle (citrate, α-ketoglutarate, aspartate, asparagine), the PPP (6-phosphogluconate, ribose-5-phosphate), and nucleotide biosynthesis (IMP, AMP, ATP, GMP, GTP, UMP, UTP, and CTP) was significantly reduced in KO cells relative to K27M in all models. We did not observe changes in the ^13^C labeling of glutamate, succinate, malate, ADP, GDP, UDP, or CDP in KO cells relative to K27M cells (Fig. 1G-1K). Loss of H3K27M expression depleted ATP, NAD+, and NADPH (Supplementary Fig. S1A-S1F) in DMG cells, consistent with the inhibition of glycolysis and the PPP.

To confirm these results using an alternate approach, we examined murine 24-B7 cells that were originally generated by brainstem-targeted intra-uterine electroporation of plasmids expressing histone H3.3 K27M, dominant negative TP53, and constitutively active PDGFRA into mice and subsequently established as neurospheres in culture (30). As controls, we examined murine 24-D3 cells that were generated by the same process but expressed wild-type histone H3.3 (30). These cells have been shown to recapitulate the histopathology, microenvironment, and therapeutic response of human DMG (30). As expected (30), 24-B7 cells showed expression of the mutant H3K27M protein, globally reduced H3K27me3, and globally elevated H3K27Ac relative to 24-D3 cells (Fig. 1L). Importantly, consistent with the results from the patient-derived models, [U-^13^C]-glucose metabolism via glycolysis (hexose phosphate, fructose-1,6-bisphosphate, 3-phosphoglycerate, lactate), the TCA cycle (citrate, α-ketoglutarate, aspartate, asparagine), the PPP (6-phosphogluconate, ribose-5-phosphate), and nucleotide (IMP, AMP, ATP, GMP, GTP, UMP, UTP, and CTP) biosynthesis was significantly higher in 24-B7 cells relative to 24-D3 (Fig. 1M). Collectively, these results indicate that the H3K27M mutation upregulates glucose metabolism via glycolysis, the TCA cycle, the PPP, and *de novo* nucleotide biosynthesis in patient-derived DMG models.

### The H3K27M mutation upregulates the rate-limiting glycolytic enzyme phosphoglycerate kinase 1 (PGK1) in patient-derived DMG cells

Given the striking reduction in lactate production induced by H3K27M silencing, we questioned whether lactate plays a functional role in DMG growth and metabolism. To this end, we first examined the effect of H3K27M silencing on the expression of glycolytic enzymes and transporters in DMG cells. As shown in Fig. 1N-1Q, the expression of PGK1, which converts 1,3-bisphosphoglycerate to 3-phosphoglycerate while producing ATP (see the schematic in Fig. 1A), was significantly reduced in KO cells relative to K27M in all DMG models. PGK1 expression was also significantly elevated in 24-B7 cells relative to 24-D3 (Fig. 1R). These results are consistent with the significant reduction in 3-phosphoglycerate, lactate, and ATP induced by H3K27M silencing in DMG cells (Fig. 1G-1K) and the reciprocal increase in these metabolites in 24-B7 cells relative to 24-D3 (Fig. 1M). Except for the monocarboxylate transporter MCT4 (encoded by *SLC16A3*; Supplementary Fig. S1G-S1I), none of the other glycolytic enzymes or transporters were consistently altered by H3K27M status in DMG models.

The H3K27M mutation is known to induce epigenetic alterations that activate gene expression including a global reduction in transcriptionally repressive histone H3K27me3 at gene promoters (3). As shown in Fig. 1S-1T, chromatin immunoprecipitation quantitative PCR (ChIP-QPCR) showed a significant increase in histone H3K27me3 at the *PGK1* promoter in KO cells relative to K27M for all DMG models. Conversely, we observed a significant decrease in histone H3K27me3 at the PGK1 promoter in 24-B7 cells relative to 24-D3 (Fig. 1U). Taken together, these studies indicate that the H3K27M mutation induces epigenetic alterations that upregulate PGK1 expression in DMGs.

### Lactate is essential for NTP synthesis in patient-derived DMG cells

To delineate the role of lactate in DMG cells, we first took the approach of depleting lactate by silencing PGK1 via CRISPRi (Fig. 2A) and then identifying metabolic and phenotypic changes that are rescued by supplementation with exogenous lactate. As shown in Fig. 2B-2F, as expected, PGK1 silencing significantly downregulated [U-^13^C]-glucose metabolism to 3-phosphoglycerate, lactate, citrate, and α-ketoglutarate in all patient-derived DMG models. Strikingly, PGK1 silencing also caused a significant reduction in the ^13^C labeling of NTPs (GTP, UTP, and CTP) except ATP in DMG cells (Fig. 2B-2F). PGK1 ablation did not alter ^13^C labeling of NDPs (GDP, UDP, CDP, ADP) in DMG cells (Fig. 2B-2F). Importantly, supplementation with exogenous lactate rescued the synthesis of GTP, UTP, and CTP from [U-^13^C]-glucose in patient-derived DMG cells (Fig. 2G-2I). This effect was specific to lactate because supplementation with exogenous citrate or cell-permeable dimethyl α-ketoglutarate did not rescue NTP levels (Fig. 2G-2I). We confirmed that PGK1 silencing also depleted the pool sizes of GTP, UTP, and CTP, an effect that was specifically rescued by lactate but not by citrate or dimethyl α-ketoglutarate in DMG cells (Supplementary Fig. S2A-S2C).

**Figure 2.**
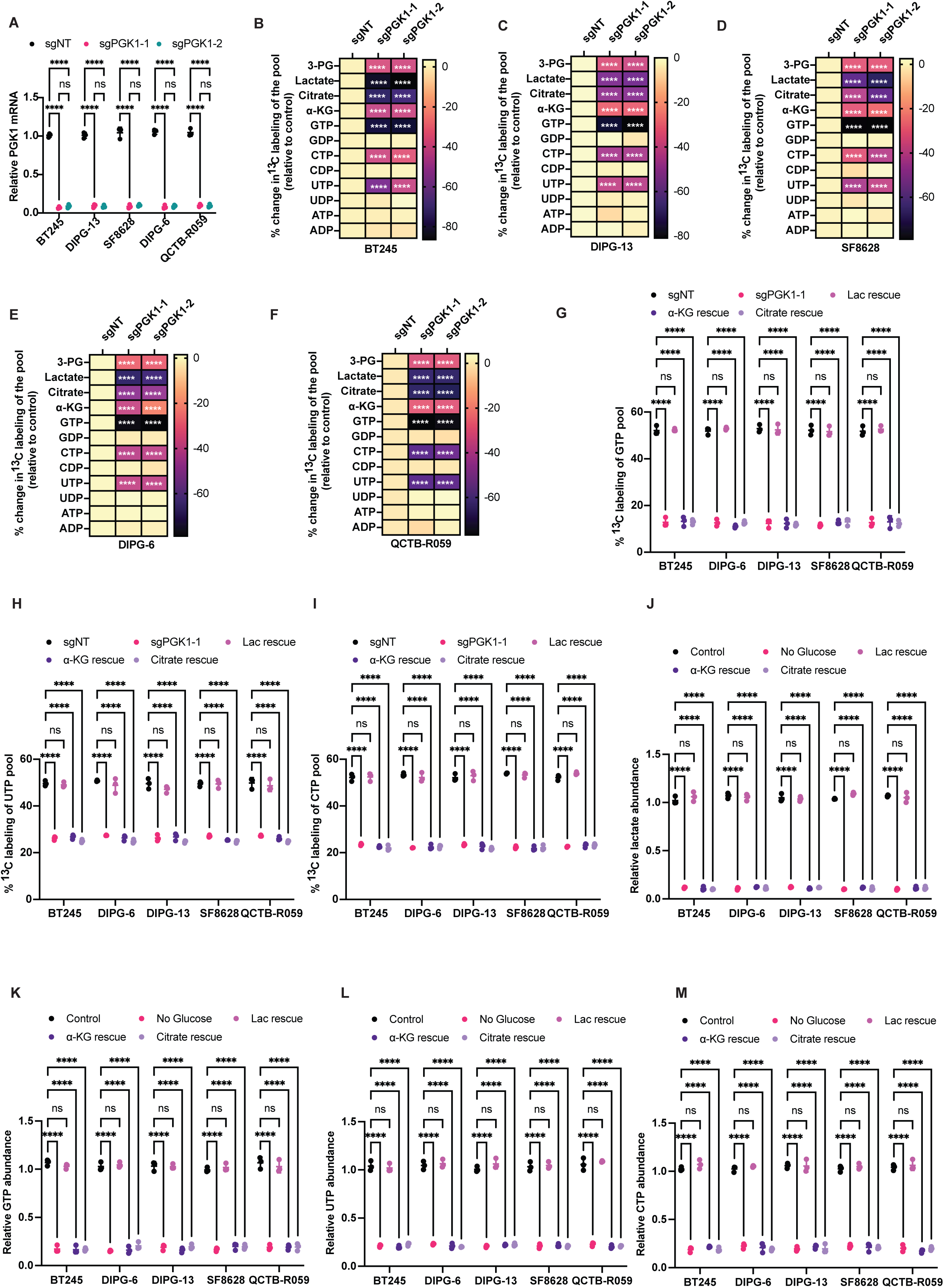
Lactate is essential for NTP synthesis in DMG cells. **(A)** Verification of loss of PGK1 mRNA in BT245, DIPG-13, SF8628, DIPG-6, and QCTB-R059 cells expressing sgRNA against PGK1 (sgPGK1-1 and sgPGK-1-2) or a non-targeting control (sgNT). Effect of PGK1 silencing on [U-^13^C]-glucose metabolism in the BT245 **(B)**, DIPG-13 **(C)**, SF8628 **(D)**, DIPG-6 **(E)**, and QCTB-R059 **(F)** models. Effect of supplementation with exogenous 1 mM sodium lactate (referred to as Lac rescue), 1 mM sodium citrate (Citrate rescue), or 500 μM dimethyl α-ketoglutarate (α-KG rescue) on the synthesis of ^13^C-labeled GTP **(G)**, UTP **(H)**, and CTP **(I)** in BT245, DIPG-13, SF8628, DIPG-6, and QCTB-R059 cells expressing sgRNA against PGK1 or a non-targeting control. **(J)** Effect of glucose starvation (referred to as Low Glucose) on lactate abundance in BT245, DIPG-13, SF8628, DIPG-6, and QCTB-R059 cells. The medium was also supplemented with 1 mM sodium lactate (Lac rescue), 1 mM sodium citrate (Citrate rescue), or 500 μM dimethyl α-ketoglutarate (α-KG rescue) to assess the ability of these metabolites to restore the lactate pool. Quantification of the abundance of GTP **(K)**, UTP **(L)**, and CTP **(M)** in BT245, DIPG-13, SF8628, DIPG-6, and QCTB-R059 cells subjected to glucose starvation with or without supplementation with exogenous 1 mM sodium lactate, 1 mM sodium citrate, or 500 μM dimethyl α-ketoglutarate.

To validate these results using an orthogonal approach, we examined the effect of depleting lactate by starving DMG cells of glucose. As shown in Fig. 2J, glucose starvation significantly reduced lactate abundance in all DMG models. Supplementation with exogenous lactate but not citrate or dimethyl α-ketoglutarate restored the lactate pool size to levels observed in non-starvation controls (Fig. 2J). Importantly, like PGK1 silencing, glucose starvation depleted GTP, UTP, and CTP in patient-derived DMG cells, an effect that was specifically rescued by exogenous lactate (Fig. 2K-2M). Taken together, these findings indicate that lactate is essential for NTP synthesis and the maintenance of NTP pool size in clinically relevant patient-derived DMG models.

### Lactate is essential for cell cycle progression and DNA replication in patient-derived DMG cells

Nucleotides are the building blocks for DNA and RNA and, as such, nucleotide depletion impairs proliferation by arresting cells in the S phase of the cell cycle (31–33). We performed flow cytometry to quantify the incorporation of 5-ethynyl-2′-deoxyuridine (EdU) into newly synthesized DNA (31). As shown in Fig. 3A-3D, both PGK1 silencing and glucose starvation arrested cells in the S phase of the cell cycle. Importantly, supplementation with lactate rescued S phase arrest in patient-derived DMG cells (Fig. 3A-3D).

**Figure 3.**
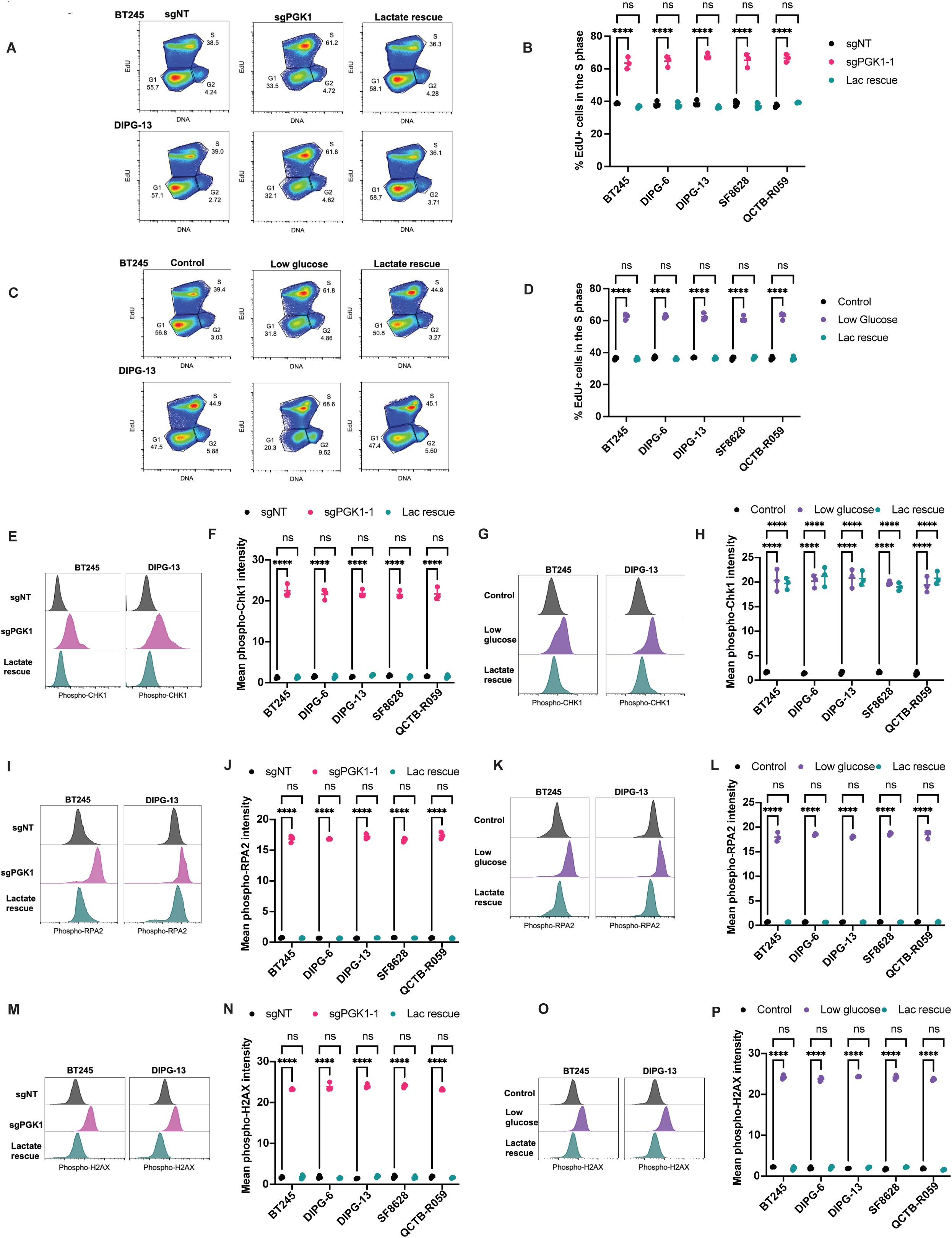
Lactate is essential for DNA replication and proliferation in DMGs. **(A)** Representative flow cytometric dot plots showing the effect of PGK1 silencing on the distribution of EdU+ cells in the G1, S, and G2/M phases for the BT245 and DIPG-13 models. **(B)** Quantification of the % of EdU+ cells in the S phase of the cell cycle in sgNT, sgPGK1, and sgPGK1 cells supplemented with 1 mM sodium lactate for the BT245, DIPG-13, SF8628, DIPG-6, and QCTB-R059 models. **(C)** Representative flow cytometric dot plots showing the effect of glucose starvation on the distribution of EdU+ cells in the G1, S, and G2/M phases for the BT245 and DIPG-13 models. **(D)** Effect of glucose starvation and lactate rescue on the % of EdU+ cells in the S phase of the cell cycle for the BT245, DIPG-13, SF8628, DIPG-6, and QCTB-R059 models. **(E)** Representative flow cytometric histograms showing the levels of phosphorylated CHK1 (S345) in sgNT, sgPGK1, and sgPGK1 cells supplemented with 1 mM sodium lactate for the BT245 and DIPG-13 models. **(F)** Quantification of the effect of PGK1 silencing and lactate rescue on the expression of phosphorylated CHK1 (S345) for the BT245, DIPG-13, SF8628, DIPG-6, and QCTB-R059 models. **(G)** Representative flow cytometric histograms showing the levels of phosphorylated CHK1 (S345) in BT245 and DIPG-13 cells subjected to glucose starvation and lactate rescue. **(H)** Quantification of the effect of glucose starvation and lactate rescue on the expression of phosphorylated CHK1 (S345) for the BT245, DIPG-13, SF8628, DIPG-6, and QCTB-R059 models. **(I)** Representative flow cytometric histograms showing the levels of phosphorylated RPA2 (S33) in sgNT, sgPGK1, and sgPGK1 cells supplemented with 1 mM sodium lactate for the BT245 and DIPG-13 models. **(J)** Quantification of the effect of PGK1 silencing and lactate rescue on the expression of phosphorylated RPA2 (S33) for the BT245, DIPG-13, SF8628, DIPG-6, and QCTB-R059 models. **(K)** Representative flow cytometric histograms showing the levels of phosphorylated RPA2 (S33) in BT245 and DIPG-13 cells subjected to glucose starvation and lactate rescue. **(L)** Quantification of the effect of glucose starvation and lactate rescue on the expression of phosphorylated RPA2 (S33) for the BT245, DIPG-13, SF8628, DIPG-6, and QCTB-R059 models. **(M)** Representative flow cytometric histograms showing the levels of phosphorylated H2AX (S139) in sgNT, sgPGK1, and sgPGK1 cells supplemented with 1 mM sodium lactate for the BT245 and DIPG-13 models. **(N)** Quantification of the effect of PGK1 silencing and lactate rescue on the expression of phosphorylated H2AX (S139) for the BT245, DIPG-13, SF8628, DIPG-6, and QCTB-R059 models. **(O)** Representative flow cytometric histograms showing the levels of phosphorylated H2AX (S139) in BT245 and DIPG-13 cells subjected to glucose starvation and lactate rescue. **(P)** Quantification of the effect of glucose starvation and lactate rescue on the expression of phosphorylated H2AX (S139) for the BT245, DIPG-13, SF8628, DIPG-6, and QCTB-R059 models.

Since DNA replication requires nucleotides, we tested whether nucleotide depletion due to PGK1 silencing triggered DNA replication stress signaling (31–33). The activation of the ATR pathway is a reliable marker for replication stress (33). Nucleotide depletion induces the binding of replication protein A2 (RPA2) to the DNA, which recruits ATR. The presence of DNA-bound RPA2 activates ATR, which phosphorylates both RPA2 at serine 33 (S33) and the downstream effector CHK1 at serine 345 (S345). As shown in Fig. 3E-3L, both PGK1 silencing and glucose starvation resulted in a significant increase in the levels of phosphorylated CHK1 (S345) and phosphorylated RPA2 (S33) in patient-derived DMG cells. Since replication stress can also lead to the onset of DNA damage (33), we tested for phosphorylation of histone H2AX at serine 139 (S139), which is a robust marker of DNA damage (33). As shown in Fig. 3M-3P, H2AX phosphorylation was higher in PGK1-silenced and glucose-starved cells relative to the corresponding controls. Importantly, replication stress and DNA damage were rescued by supplementation with exogenous lactate in our DMG models (Fig. 3E-3P). Collectively, these results identify a functional role for lactate in facilitating nucleotide biosynthesis and tumor proliferation in patient-derived DMG models.

### Lactate induces lysine lactylation of the nucleoside diphosphate kinase NME1 in patient-derived DMG cells

Next, we examined the mechanism by which lactate upregulates *de novo* NTP biosynthesis in DMGs. Since our data indicated that both PGK1 silencing and glucose starvation reduced ^13^C labeling and pool sizes of NTPs (GTP, UTP, CTP) but not of the corresponding NDPs (GDP, UDP, CDP), we quantified nucleoside diphosphate kinase (NDPK) activity in our models. The NDPK enzymes, encoded by the NME family of genes *NME1-4*, catalyze the transfer of the γ-phosphate from ATP to an NDP (GDP, UDP, or CDP) to form the corresponding NTP (GTP, UTP, and CTP; see Fig. 4A) (34). As shown in Fig. 4B-4C, NDPK activity was significantly reduced in PGK1-silenced or glucose-starved cells vs. the corresponding controls in all patient-derived DMG models. Importantly, supplementation with lactate rescued NDPK activity, indicating that lactate is critical for NDPK activity in DMG cells (Fig. 4B-4C).

**Figure 4.**
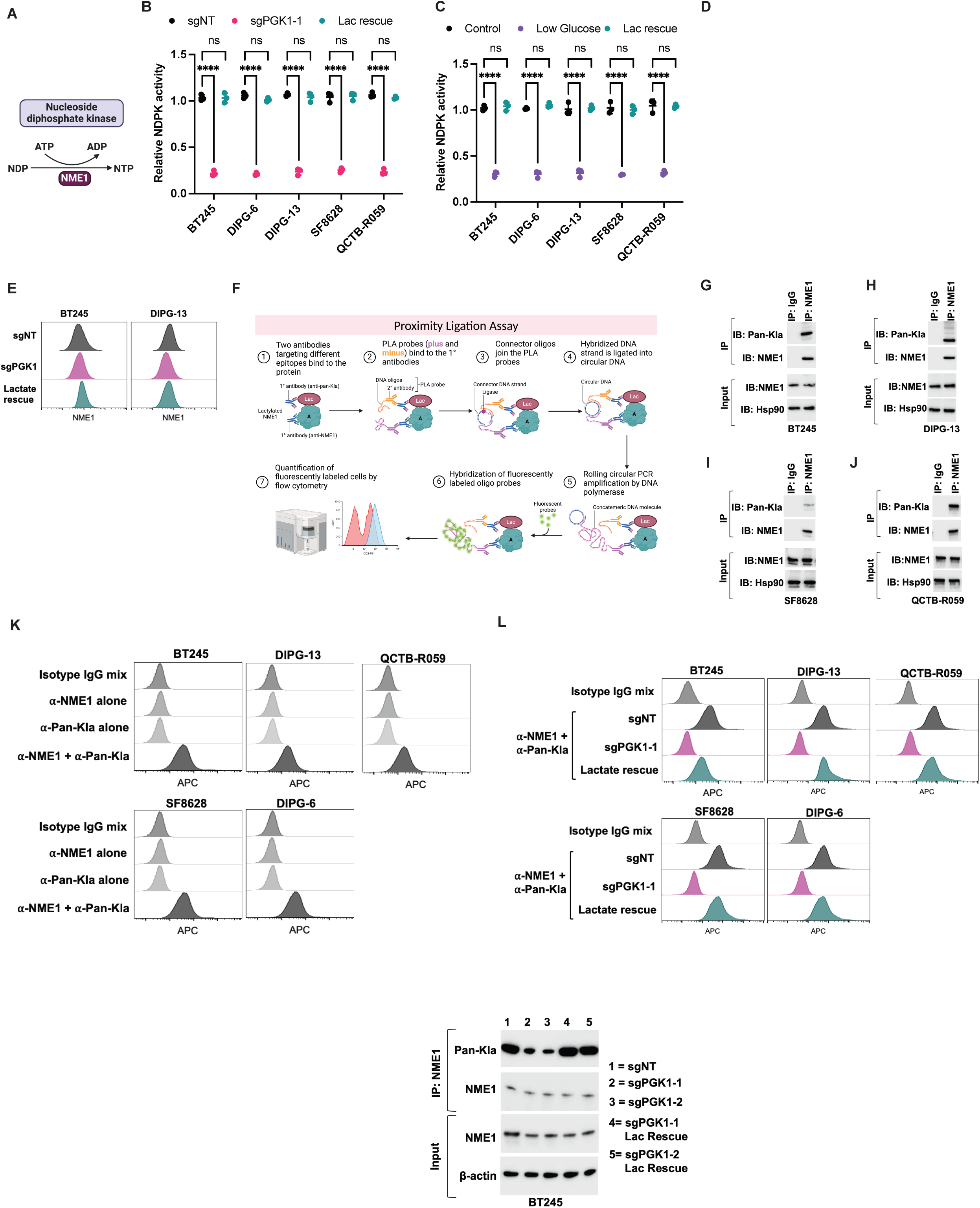
Lactate induces lactylation of NME1 in DMG cells. **(A)** Schematic illustration of the activity of NME1 in catalyzing the transfer of the γ-phosphate from ATP to an NDP to produce the corresponding NTP. The figure was created using Biorender. Effect of PGK1 silencing **(B)** or glucose starvation **(C)** on NDPK activity in the BT245, DIPG-13, SF8628, DIPG-6, and QCTB-R059 models. Cells were supplemented with 1 mM sodium lactate to assess the ability of lactate to restore NDPK activity. **(D)** Effect of silencing NME1, NME2, NME3, or NME4 on the ability of lactate to rescue NDPK activity in BT245 cells subjected to glucose starvation. **(E)** Top panel: Representative flow cytometric histograms showing NME1 expression in sgNT, sgPGK1, and sgPGK1 cells supplemented with 1 mM sodium lactate for the BT245 and DIPG-13 models. Bottom panel: Representative flow cytometric histograms showing NME1 expression in BT245 and DIPG-13 cells subjected to glucose starvation and lactate rescue. **(F)** Schematic illustration of the principle of the PLA. Briefly, cells are fixed and treated with 2 antibodies raised in different species that target different epitopes on NME1 (e.g., mouse anti-NME1 and rabbit anti-Pan-Kla). Secondary antibodies conjugated with oligonucleotides (PLA probe MINUS and PLA probe PLUS) are then added to the samples. This is followed by a ligation step in which connector DNA oligonucleotides and a ligase are added. The DNA oligonucleotides hybridize with the PLA probes, and the ligase will form a rolling circle if the 2 antibodies are in proximity to each other (<40 nm apart). An amplification solution consisting of nucleotides, a DNA polymerase, and fluorescently labeled oligonucleotides is then added. The DNA polymerase amplifies the circular DNA probe via rolling circle amplification, and the fluorescently labeled oligonucleotides hybridize to a specific sequence in the amplified DNA to yield a fluorescence signal that is readily detectable by flow cytometry. The figure was created using Biorender. Western blots demonstrating NME1 lactylation in the BT245 **(G)**, DIPG-13 **(H)**, SF8628 **(I)**, and QCTB-R059 **(J)** models. Lysates were immunoprecipitated with an anti-NME1 antibody or isotype IgG control and probed for lactylation using an anti-Pan-Kla antibody. IP refers to immunoprecipitation and IB refers to immunoblotting. The top panel shows the IP, and the bottom panel shows the input for IB at 10%. **(K)** Representative flow cytometric histograms of the PLA in the BT245, DIPG-13, SF8628, DIPG-6, and QCTB-R059 models. Cells were treated with a mixture of the mouse anti-NME1 and rabbit Pan-Kla antibodies (referred to as α-NME1 + α-Pan-Kla). Cells treated with the mixture of the corresponding isotype IgG antibodies (referred to as Isotype IgG mix) or with either primary antibody alone (referred to as α-NME1 alone or α-Pan-Kla alone) were used as negative controls. **(L)** Representative flow cytometric histograms of the PLA in sgNT, sgPGK1, and sgPGK1 cells supplemented with 1 mM sodium lactate for the BT245, DIPG-13, SF8628, DIPG-6, and QCTB-R059 models. Cells were treated with a mixture of the mouse anti-NME1 and rabbit Pan-Kla antibodies. Cells treated with the mixture of the corresponding isotype IgG were used as negative controls. **(M)** Representative flow cytometric histograms of the PLA in BT245, DIPG-13, SF8628, and DIPG-6 cells subjected to glucose starvation or lactate rescue. Cells were treated with a mixture of the mouse anti-NME1 and rabbit Pan-Kla antibodies. Cells treated with the mixture of the corresponding isotype IgG were used as negative controls. Western blots demonstrating the effect of PGK1 silencing and lactate rescue on NME1 lactylation in the BT245 **(N)** and DIPG-13 **(O)** models. Lysates were immunoprecipitated with an anti-NME1 antibody and probed for lactylation using an anti-Pan-Kla antibody. The top panel shows the IP, and the bottom panel shows the input for IB at 10%.

NDPK is encoded by the NME family of genes *NME1-4* (34). To identify the NDPK family member that was modulated by lactate, we systematically examined the effect of silencing NME1-4 on the ability of lactate to rescue NDPK activity in glucose-starved DMG cells. As shown in Fig. 4D and Supplementary Fig. S3A, silencing NME1 but not NME2, NME3, or NME4 abrogated the ability of lactate to rescue NDPK activity in BT245 and DIPG-13 cells subjected to glucose starvation. These results suggest that lactate drives NME1-based NDPK activity in patient-derived DMG models.

We then examined the mechanism by which lactate activates NME1. Lactate can activate gene expression via lactylation of histones, most commonly histone H3K18 lactylation (19,20). However, we did not observe changes in NME1 mRNA (Supplementary Fig. S3B-S3C), NME1 protein (Fig. 4E and Supplementary Fig. S3D-S3E), or the enrichment of the H3K18 lactylation mark at the NME1 promoter (Supplementary Fig. S3F-S3G) following PGK1 silencing or glucose starvation in patient-derived DMG cells.

Another mechanism by which lactate can influence protein function is by covalent modification at lysine residues (19,20). We used a proximity ligation assay (PLA) coupled with flow cytometry to examine whether NME1 is being lactylated in DMG cells (see schematic illustration in Fig. 4F). The PLA enables *in situ* detection of endogenous protein modifications with high specificity and sensitivity (35–37). The assay uses two primary antibodies raised in different species i.e., rabbit and mouse, to detect two unique targets, e.g., NME1 and the pan-lactyl lysine (Pan-Kla) moiety. A pair of oligonucleotide-labeled secondary antibodies (PLA probes) then binds to the primary antibodies. Next, hybridizing connector oligonucleotides join the PLA probes only if they are in close proximity to each other and the ligase forms a closed, circle DNA template that is required for rolling-circle amplification. The PLA probe acts as a primer for DNA polymerase, which generates an amplified fluorescent signal that can be visualized by flow cytometry (35–37) (Fig. 4F). In comparison to traditional methods like immunoprecipitation that are suited for high abundance proteins and require large sample volumes, the PLA allows high-throughput visualization of endogenous low-abundance proteins in as little as ∼100,000 cells from cell lines, xenograft tissue, or patient biopsies (35–37). Of note, the PLA allows for quantification of the extent of protein modification across different samples. To validate the PLA for the detection of NME1 lactylation, we first examined NME1 lactylation by immunoprecipitation followed by immunoblotting in patient-derived DMG cells. We immunoprecipitated NME1 from cell lysates using an anti-NME1 antibody and then blotted the immunoprecipitate with a Pan-Kla antibody. Immunoprecipitation of cell lysates with isotype IgG was used as a control. As shown in Fig. 4G-4J, we observed NME1 lactylation in DMG lysates immunoprecipitated with an anti-NME1 antibody and subsequently blotted using a Pan-Kla antibody. Importantly, the PLA with the combination of the anti-NME1 and anti-Pan-Kla antibodies showed a strong fluorescent signal indicating lactylation of NME1 in all patient-derived DMG models (Fig. 4K and Supplementary Fig. S3H). We confirmed that incubation of DMG cells with a mixture of isotype IgGs, anti-NME1 antibody alone, or anti-pan-lactyl lysine antibody alone did not yield fluorescence (Fig. 4K and Supplementary Fig. S3H).

We then examined the effect of depleting lactate by silencing PGK1 or glucose starvation on NME1 lactylation in DMG cells. As shown in Fig. 4L-4M and Supplementary Fig. S3I-S3J, both PGK1 silencing and glucose starvation significantly reduced NME1 lactylation in DMG cells, an effect that was rescued by supplementation with exogenous lactate. We also confirmed the validity of these results by immunoprecipitation followed by immunoblotting. As shown in Fig. 4N-4O and Supplementary Fig. S3K, PGK1 silencing abrogated NME1 lactylation in BT245 and DIPG-13 cells, an effect that was rescued by supplementation with exogenous lactate. Collectively, the results presented here indicate that lactate covalently modifies NME1 via lysine lactylation in patient-derived DMG cells.

### Lactate activates NME1 via lactylation of lysine 49 in patient-derived DMG cells

To further confirm the role of lactylation in activating NME1, we sought to identify the lysine residue in NME1 that is lactylated. NME1 (protein accession #: P15531) has 10 lysine residues. We systematically examined the effect of expressing either wild-type or mutant NME1 in which each lysine is replaced with an arginine, which prevents lactylation (K12R, K26R, K31R, K39R, K49R, K56R, K66R, K85R, K100R, K128R) in BT245 cells. As shown in the PLA in Fig. 5A and Supplementary Fig. S4A, the K49R mutation abrogated NME1 lactylation in BT245 cells, returning it to levels observed in the negative isotype IgG controls. We also confirmed these results by immunoprecipitation followed by immunoblotting in BT245 cells expressing wild-type or K49R mutant NME1 (Fig. 5B). Importantly, the K49R mutation significantly reduced NDPK activity and selectively depleted GTP, UTP, and CTP, leading to S-phase arrest and a significant increase in doubling time (Fig. 5C-5G). Consistent with nucleotide depletion, the K49R mutation activated replication stress and DNA damage as indicated by elevated phosphorylation of CHK1 (S345), RPA2 (S33), and H2AX (S139) in K49R mutant cells relative to wild-type (Fig. 5H-5M). Collectively, our results indicate that lactylation at K49 activates NME1, thereby upregulating NTP biosynthesis and driving tumor proliferation in DMGs.

**Figure 5.**
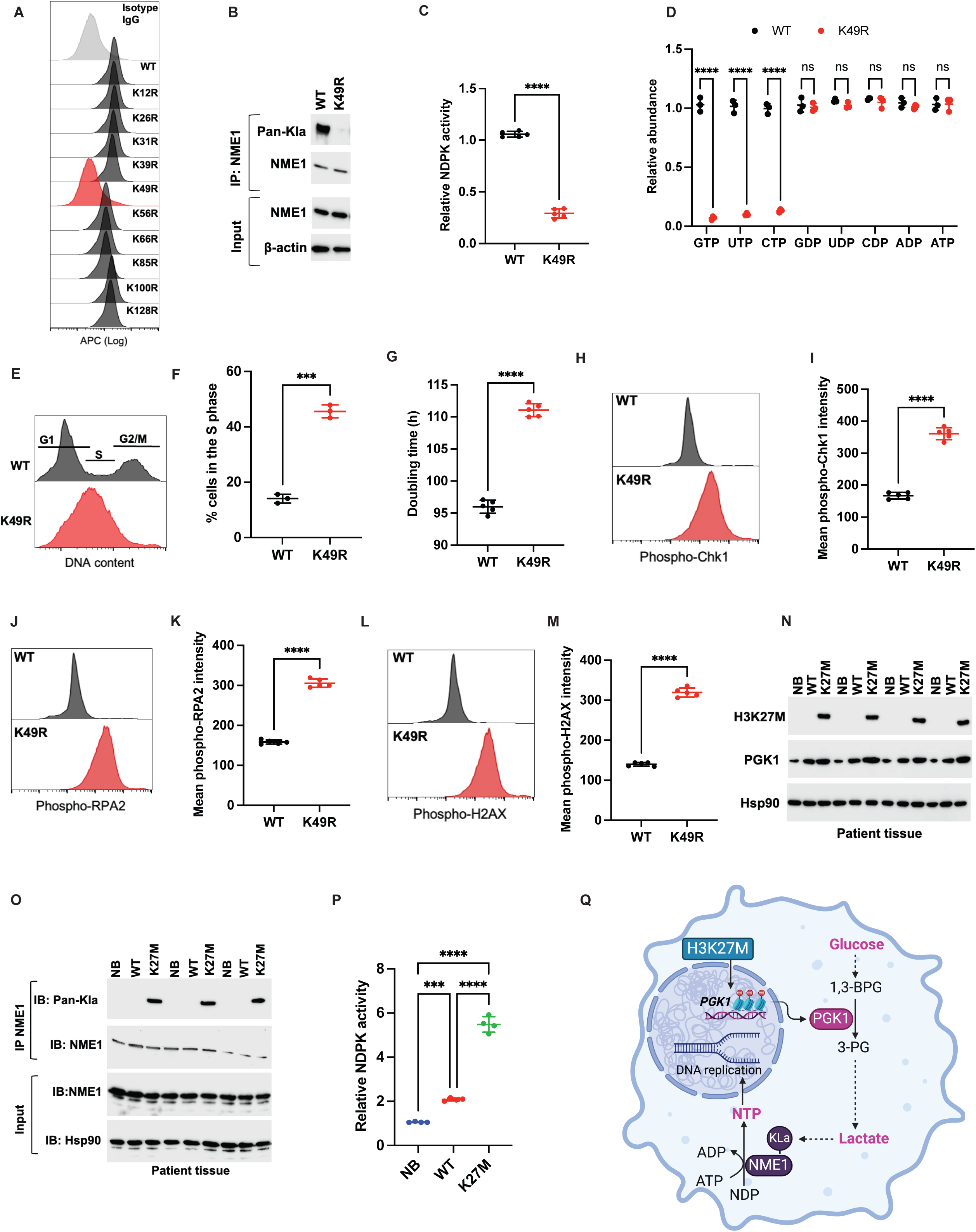
Lactate causes NME1 lactylation at K49. **(A)** Representative flow cytometric histograms of the PLA in BT245 cells expressing wild-type (WT) NME1 or NME1 in which each lysine residue is mutated to arginine. Cells were treated with a mixture of the mouse anti-NME1 and rabbit Pan-Kla antibodies. Cells treated with the mixture of the corresponding isotype IgG were used as negative controls. **(B)** Western blots of NME1 lactylation in BT245 cells expressing WT or K49R mutant NME1. Lysates were immunoprecipitated with an anti-NME1 antibody and then immunoblotted with the anti-Pan-Kla antibody. The top panel shows the IP, and the bottom panel shows the input for IB at 10%. Effect of the K49R mutation on NDPK activity **(C)** and NTP abundance **(D)** in BT245 cells. **(E)** Representative flow cytometric histograms of cell cycle distribution in BT245 cells expressing WT or K49R mutant NME1. **(F)** Quantification of the % of cells in the S phase of the cell cycle in BT245 cells expressing WT or K49R mutant NME1. **(G)** Effect of the K49R mutation on the doubling time of BT245 cells. **(H)** Representative flow cytometric histograms showing the levels of phosphorylated CHK1 (S345) in BT245 cells expressing WT or K49R mutant NME1. **(I)** Quantification of phosphorylated CHK1 (S345) expression in BT245 cells expressing WT or K49R mutant NME1. **(J)** Representative flow cytometric histograms showing the levels of phosphorylated RPA2 (S33) in BT245 cells expressing WT or K49R mutant NME1. **(K)** Quantification of phosphorylated RPA2 (S33) expression in BT245 cells expressing WT or K49R mutant NME1. **(L)** Representative flow cytometric histograms showing the levels of phosphorylated H2AX (S139) in BT245 cells expressing WT or K49R mutant NME1. **(M)** Quantification of phosphorylated H2AX (S139) expression in BT245 cells expressing WT or K49R mutant NME1. **(N)** Western blots showing the expression of mutant H3K27M and PGK1 in biopsies from patients with H3K27M mutant (K27M) or H3 wild-type (WT) tumors. Normal pontine brain tissue (NB) was used as an added control. **(O)** Western blots demonstrating NME1 lactylation in biopsies from patients with H3K27M mutant (K27M), H3 wild-type (WT) tumors, or normal brain tissue (NB). Lysates were immunoprecipitated with an anti-NME1 antibody and probed for lactylation using an anti-Pan-Kla antibody. The top panel shows the IP, and the bottom panel shows the input for IB at 10%. **(P)** NDPK activity in biopsies from patients with H3K27M mutant (K27M), H3 wild-type (WT) tumors, or normal brain tissue (NB). **(Q)** Schematic summary of the mechanism by which the H3K27M mutation drives NTP synthesis and tumor proliferation in DMGs. The H3K27M mutation upregulates PGK1 expression via reduced H3K27me3 enrichment at the PGK1 promoter. Elevated PGK1 drives the glycolytic production of lactate, which, in turn, activates NME1 via lactylation at K49. Upregulated NME1 facilitates the synthesis of NTPs needed for DNA replication and tumor proliferation in DMGs.

### NME1 lactylation is significantly upregulated in H3K27M patient tumors relative to H3 wild-type tumors

To confirm the clinical relevance of our studies, we examined biopsies obtained from patients with H3K27M or H3 wild-type tumors. As added controls, we examined normal brain tissue obtained at autopsy. We confirmed the presence of the H3K27M mutation in H3K27M biopsies relative to H3 wild-type and normal brain (Fig. 5N). As shown in Fig. 5N, PGK1 expression was significantly higher in H3K27M patient biopsies but not H3 wild-type biopsies and normal brains. Importantly, NME1 lactylation was observed specifically in H3K27M patient biopsies but not H3 wild-type biopsies or normal brains (Fig. 5O). We also confirmed that NDPK activity (Fig. 5P) and NTP abundance (Supplementary Fig. S4B) were significantly higher in H3K27M biopsies relative to H3 wild-type and normal brains. Taken together, these results validate the relevance of our preclinical findings to DMG patients and identify a unique H3K27M-lactate-NME1 cascade that promotes NTP synthesis and tumor proliferation in DMGs (see schematic summary in Fig. 5Q).

### DMI-detectable lactate production from [6,6-^2^H]-glucose provides a readout of the H3K27M mutation in patient-derived DMG cells

Given the mechanistic association between the H3K27M mutation, lactate, and tumor proliferation (see Fig. 5Q), we questioned whether tracing [6,6-^2^H]-glucose metabolism to lactate using DMI would provide a non-invasive readout of the H3K27M mutation in DMG cells. To this end, we labeled K27M and KO cells with [6,6-^2^H]-glucose and quantified lactate production in cell suspensions for the BT245, SF8628, DIPG-6, DIPG-13, and QCTB-R059 models. As shown in the representative ^2^H-MR spectra in Fig. 6A, we observed the production of [3,3’-^2^H]-lactate (1.3 ppm) from [6,6’-^2^H]-glucose (3.75 ppm) in K27M cells. Importantly, silencing the H3K27M mutation significantly reduced [3,3’-^2^H]-lactate production from [6,6-^2^H]-glucose in all patient-derived DMG models (Fig. 6A-6C). We did not observe a significant difference in the production of ^2^H-glx (summed resonance from glutamate and glutamine at 2.4 ppm) in K27M or KO cells (Supplementary Fig. S5A-S5C), consistent with the observation that silencing the H3K27M mutation did not alter ^13^C-labeling of glutamate from [U-^13^C]-glucose in DMG cells (see Fig. 1G-1K). Of note, we confirmed that [3,3’-^2^H]-lactate production from [6,6-^2^H]-glucose was significantly higher in the isogenic H3K27M 24-B7 cells relative to H3 wild-type 24-D3 cells (Fig. 6D). Taken together, these results indicate that [3,3’-^2^H]-lactate production from [6,6-^2^H]-glucose is a quantitative biomarker of the H3K27M mutation in clinically relevant DMG cells.

**Figure 6.**
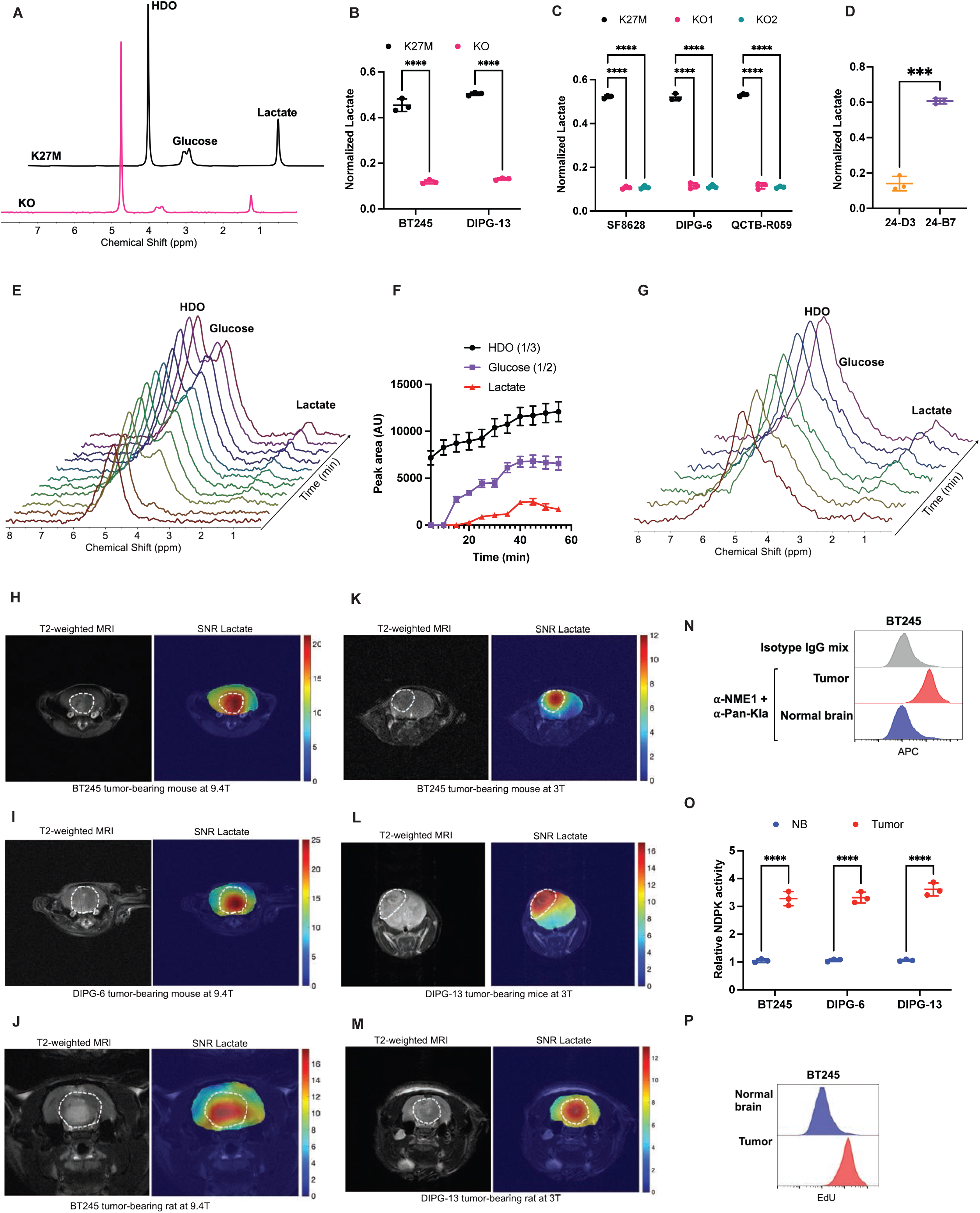
Spatially mapping [6,6’-^2^H]-glucose metabolism to lactate enables visualization of the metabolically active tumor lesion in preclinical DMG models. **(A)** Representative ^2^H-MR spectra from BT245 K27M and KO cells incubated in media containing 25 mM [6,6’-^2^H]-glucose for 72 h. **(B)** Quantification of [3,3’-^2^H]-lactate production from [6,6’-^2^H]-glucose in K27M and KO cells for the BT245 and DIPG-13 models. **(C)** Quantification of [3,3’-^2^H]-lactate production from [6,6’-^2^H]-glucose in K27M and KO cells for the SF8628, DIPG-6, and QCTB-R059 models. **(D)** Quantification of [3,3’-^2^H]-lactate production from [6,6’-^2^H]-glucose in isogenic 24-B7 (H3K27M) and 24-D3 (H3 wild-type) cells. **(E)** Representative spectral array showing the dynamic production of [3,3’-^2^H]-lactate from [6,6’-^2^H]-glucose in a mouse bearing an intracranial DIPG-6 tumor implanted in the pons. Data was acquired using a non-localized sequence on a 9.4T scanner after intravenous administration of [6,6’-^2^H]-glucose. **(F)** Quantification of the kinetics of HDO, glucose, and lactate following intravenous administration of [6,6’-^2^H]-glucose into mice bearing intracranial pontine DIPG-6 tumors at 9.4T. **(G)** Representative spectral array showing the dynamic production of [3,3’-^2^H]-lactate from [6,6’-^2^H]-glucose in a mouse bearing an intracranial BT245 tumor implanted in the cortex. Data was acquired using a non-localized sequence on a 3T scanner after intravenous administration of [6,6’-^2^H]-glucose. **(H-J)** Representative 2D CSI data acquired on a 9.4T scanner after intravenous administration of [6,6’-^2^H]-glucose into a mouse bearing an intracranial BT245 tumor implanted in the pons **(H)**, a mouse with an intracranial DIPG-6 tumor implanted in the pons **(I)**, or a rat with an intracranial BT245 tumor implanted in the pons **(J)**. In each case, the left panel shows the T2-weighted MRI with the tumor outlined in white and the right panel shows the corresponding heatmap of the SNR of [3,3’-^2^H]-lactate. **(K-M)** Representative 2D CSI data acquired on a 3T scanner after administration of [6,6’-^2^H]-glucose into a mouse bearing an intracranial BT245 tumor implanted in the cortex **(K)**, a mouse with an intracranial DIPG-13 tumor implanted in the cortex **(L)**, or a rat with an intracranial DIPG-13 tumor implanted in the pons **(M)**. In each case, the left panel shows the T2-weighted MRI with the tumor outlined in white and the right panel shows the corresponding heatmap of the SNR of [3,3’-^2^H]-lactate. **(N)** Representative flow cytometric histograms of the PLA in tumor and contralateral normal brain tissue resected from a mouse bearing an intracranial BT245 tumor after the acquisition of DMI data. Cells were treated with a mixture of the mouse anti-NME1 and rabbit Pan-Kla antibodies. Cells treated with the mixture of the corresponding isotype IgG were used as negative controls. **(O)** Quantification of NDPK activity in tumor and contralateral normal brain tissue resected from mice bearing intracranial BT245, DIPG-13, or DIPG6 tumors. **(P)** Representative flow cytometric histograms of EdU incorporation into tumor and contralateral normal brain tissue resected from a mouse bearing an intracranial BT245 tumor after the acquisition of DMI data.

### Spatially mapping lactate production from [6,6-^2^H]-glucose enables visualization of the metabolically active tumor lesion in mice and rats bearing orthotopic DMG xenografts *in vivo*

Next, we questioned whether [6,6-^2^H]-glucose can be used to visualize tumor burden in preclinical DMG models *in vivo.* To this end, we first acquired non-localized ^2^H-MR spectra from the brain after intravenous injection of [6,6’-^2^H]-glucose into mice bearing intracranial DIPG-6 tumors at a magnetic field strength of 9.4T. As shown in the representative spectral array in Fig. 6E and the quantification in Fig. 6F, the peak for semi-heavy water (HDO; 4.75 ppm) was the only signal observed prior to intravenous injection of [6,6’-^2^H]-glucose into DIPG-6 tumor-bearing mice. Following administration of [6,6’-^2^H]-glucose, a peak for [6,6’-^2^H]-glucose at 3.75 ppm was observed followed by the dynamic build-up of [3,3’-^2^H]-lactate at 1.3 ppm (Fig. 6E-6F). The signal-to-noise ratio (SNR) of ^2^H-glx (2.4 ppm) was not sufficient for reliable detection in DIPG-6 tumor-bearing mice *in vivo.* Importantly, we confirmed that [3,3’-^2^H]-lactate production from [6,6’-^2^H]-glucose could be quantified in mice bearing intracranial BT245 xenografts at the clinically relevant magnetic field strength of 3T (see the representative spectral array in Fig. 6G and the quantification in Supplementary Fig. S5D).

To further validate these results, we used 2D chemical shift imaging (CSI) to quantify the spatial distribution of lactate production with a spatial resolution of 70.3 μL in mice or rats bearing orthotopic patient-derived DMG xenografts at 9.4T. Visualization of the data in the form of metabolic heatmaps showed that [3,3’-^2^H]-lactate produced from [6,6-^2^H]-glucose was spatially localized to the tumor relative to the surrounding normal brain in mice bearing intracranial BT245 tumors (Fig. 6H), in mice bearing intracranial DIPG-6 tumors (Fig. 6I), and in rats bearing intracranial BT245 tumors (Fig. 6J) at 9.4T. Importantly, 2D CSI performed with a spatial resolution of 113.5 μL at clinical magnetic field strength (3T) confirmed the ability of [3,3’-^2^H]-lactate to delineate tumor burden in mice bearing intracranial BT245 tumors (Fig. 6K), in mice bearing intracranial DIPG-13 tumors (Fig. 6L), and in rats bearing intracranial DIPG-13 tumors (Fig. 6M). Quantification of the data in multiple tumor-bearing animals confirmed that the SNR of [3,3’-^2^H]-lactate was significantly higher in the tumor vs. normal brain in patient-derived DMG models *in vivo* (Supplementary Fig. S5E-S5F).

To mechanistically link ^2^H-lactate production with DMG proliferation *in vivo*, we examined tumor and normal brain tissue resected from mice bearing intracranial BT245, DIPG-13, or DIPG-6 tumors after the DMI scan. Consistent with our findings in DMG cells and patient biopsies, NME1 lactylation (Fig. 6N and Supplementary Fig. S5G), NDPK activity (Fig. 6O), and EdU incorporation (Fig. 6P and Supplementary Fig. S5H) were significantly higher in tumor vs. contralateral normal brain tissue for all models. Taken together, these results indicate that tracing ^2^H-lactate production from [6,6’-^2^H]-glucose using DMI enables non-invasive imaging of tumor burden in preclinical DMG models *in vivo*.

### DMI-detectable lactate production from [6,6-^2^H]-glucose provides a readout of response to standard-of-care and experimental therapies in patient-derived DMG cells

Accurately quantifying tumor response to therapy remains a serious challenge in the clinical management of DMG patients (13,14). Given that our results above indicated that H3K27M-driven lactylation facilitates tumor proliferation and that [6,6-^2^H]-glucose provides a readout of the H3K27M mutation, we questioned whether [6,6-^2^H]-glucose can be used to interrogate response to therapy in DMG cells. Since radiation is the standard of care for DMG patients (1,2), we quantified the effect of treatment with radiation on [6,6-^2^H]-glucose metabolism in our patient-derived DMG models. First, we established that a single dose of 10 Gy radiation significantly reduced the viability of BT245 and DIPG-13 cells (Supplementary Fig. S6A). Importantly, as shown in the representative ^2^H-MR spectra in Fig. 7A and the quantification in Fig. 7B, radiation significantly reduced [3,3’-^2^H]-lactate production relative to controls in both BT245 and DIPG-13 models.

**Figure 7.**
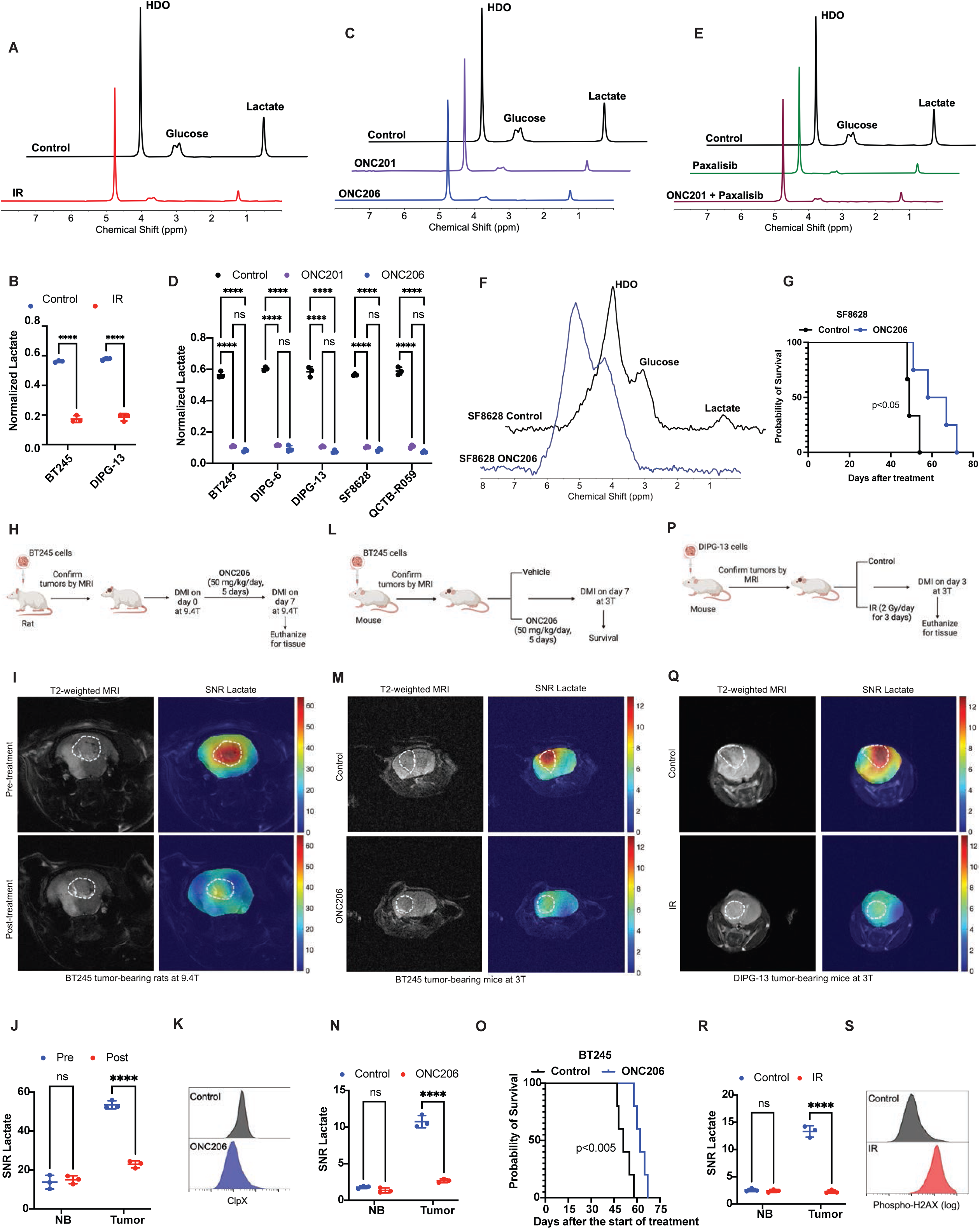
Lactate is a quantitative imaging biomarker of response to therapy in preclinical DMG models. **(A)** Representative ^2^H-MR spectra from BT245 cells that were untreated (control) or subjected to a single dose of 10 Gy radiation (IR) and concurrently incubated in media containing 25 mM [6,6’-^2^H]-glucose for 72 h. **(B)** Quantification of the effect of radiation on [3,3’-^2^H]-lactate production from [6,6’-^2^H]-glucose in BT245 and DIPG-13 cells. **(C)** Representative ^2^H-MR spectra from BT245 cells treated with vehicle, 10 μM ONC201, or 500 nM ONC206 for 72 h. Cells were concurrently incubated in media containing 25 mM [6,6’-^2^H]-glucose. **(D)** Quantification of the effect of ONC201 and ONC206 on [3,3’-^2^H]-lactate production from [6,6’-^2^H]-glucose in BT245, DIPG-13, SF8628, DIPG-6, and QCTB-R059 cells. **(E)** Representative ^2^H-MR spectra from BT245 cells treated with vehicle, 1 μM paxalisib, or the combination of 1 μM paxalisib and 5 μM ONC201 for 72 h. Cells were concurrently incubated in media containing 25 mM [6,6’-^2^H]-glucose. **(F)** Representative summed ^2^H-MR spectra from mice bearing intracranial SF8628 tumors implanted in the cortex. Mice were treated with vehicle or ONC206 for 7 days, and ^2^H-MR data was acquired by non-localized spectroscopy on a 3T scanner. **(G)** Effect of treatment with ONC206 on the survival of mice with cortical SF8628 tumors. **(H)** Schematic representation of the study design with rats bearing intracranial BT245 tumors in the pons. Once tumors were ∼80 mm^3^, 2D CSI data was acquired after intravenous administration of [6,6’-^2^H]-glucose on a 9.4T scanner. Rats were then treated with ONC206 and 2D CSI data was acquired again at day 7±1. After the DMI scan, rats were euthanized, and tissue was collected for analysis. **(I)** Representative 2D CSI data from BT245 rats pre- (top panel) and post- (bottom panel) treatment with ONC206 as described in panel H. In each case, the left panel shows the T2-weighted MRI with the tumor outlined in white and the right panel shows the corresponding heatmap of the SNR of [3,3’-^2^H]-lactate. **(J)** Quantification of the SNR of [3,3’-^2^H]-lactate in BT245 tumor-bearing rats treated with ONC206 as described in panel H. **(K)** Representative flow cytometry histograms of ClpP expression in tumor tissue resected from BT245 tumor-bearing rats treated with ONC206 as described in panel H. Tumor tissue resected from a separate cohort of vehicle-treated rats was used as control. **(L)** Schematic representation of the study design with mice bearing intracranial BT245 tumors in the cortex. Once tumors were ∼35 mm^3^, mice were treated with vehicle or ONC206, and 2D CSI data was acquired at day 7 after intravenous administration of [6,6’-^2^H]-glucose on a 3T scanner. Mice were treated longitudinally, and their survival was monitored. **(M)** Representative 2D CSI data from mice bearing cortical BT245 tumors treated with vehicle (top panel) or ONC206 (bottom panel) as described in panel L. In each case, the left panel shows the T2-weighted MRI with the tumor outlined in white and the right panel shows the corresponding heatmap of the SNR of [3,3’-^2^H]-lactate. **(N)** Quantification of the SNR of [3,3’-^2^H]-lactate in BT245 mice treated with vehicle or ONC206 as described in panel L. **(O)** Effect of treatment with ONC206 on the survival of mice with cortical BT245 tumors. **(P)** Schematic representation of the study design with mice bearing cortical DIPG-13 tumors. Once tumors were ∼35 mm^3^, mice were untreated (control) or subjected to radiation (IR) for 3 days. 2D CSI data was acquired on day 3 after intravenous administration of [6,6’-^2^H]-glucose on a 3T scanner. After the DMI scan, mice were euthanized, and tissue was collected for analysis. **(Q)** Representative 2D CSI data from the control (top panel) or IR-treated (bottom panel) DIPG-13 mice. In each case, the left panel shows the T2-weighted MRI with the tumor outlined in white and the right panel shows the corresponding heatmap of the SNR of [3,3’-^2^H]-lactate. **(R)** The SNR of [3,3’-^2^H]-lactate in DIPG-13 control or IR-treated mice. **(S)** Representative flow cytometry histograms of phospho-H2AX expression in tumor tissue resected from DIPG-13 control or IR-treated mice.

We then examined whether DMI has the potential to track response to emerging targeted therapies. The imipridones ONC201 and ONC206 are experimental therapies that have shown promising efficacy in preclinical DMG models and in clinical studies in DMG patients (6–12). Paxalisib is a brain penetrant inhibitor of the PI3K/Akt pathway that synergistically enhances response to ONC201 and is currently in clinical trials in DMG patients (10). Consistent with prior studies, we confirmed that treatment with ONC201 at a concentration of ∼10 μM and ONC206 at a concentration of ∼500 nM significantly inhibited the viability of BT245, SF8628, DIPG-6, DIPG-13, and QCTB-R059 cells (Supplementary Fig. S6B). Similarly, in line with previous studies (10), we established that treatment with paxalisib at a concentration of 1 μM or with the combination of 1 μM paxalisib and 5 μM ONC201 significantly inhibited the viability of DIPG-6 and DIPG-13 cells (Supplementary Fig. S6C). Importantly, as shown in the representative ^2^H-MR spectra in Fig. 7C and 7E, and the quantification in Fig. 7D and Supplementary Fig. S6D, [3,3’-^2^H]-lactate production from [6,6-^2^H]-glucose was significantly reduced in DMG cells treated with ONC201, ONC206, paxalisib, or the combination of paxalisib with ONC201 relative to the corresponding controls. Taken together, these results indicate that tracing ^2^H-lactate production from [6,6-^2^H]-glucose has the potential to report on the response of clinically relevant patient-derived DMG cells to the standard-of-care radiation and emerging experimental therapies.

### Lactate production from [6,6-^2^H]-glucose is a quantitative imaging biomarker of early response to radiation and targeted therapy in preclinical DMG models *in vivo*

Next, we investigated whether [6,6-^2^H]-glucose provides an early readout of DMG response to therapy *in vivo.* To this end, we first treated mice bearing intracranial patient-derived SF8628 xenografts with vehicle or ONC206 (25 mg/kg, twice daily, 5 days/week), quantified [6,6-^2^H]-glucose metabolism using non-localized ^2^H-MR spectroscopy at day 7 on a 3T scanner, and then longitudinally monitored animal survival. As shown in the representative spectra in Fig. 7F and the quantification in Supplementary Fig. S6E, [3,3’-^2^H]-lactate production from [6,6-^2^H]-glucose was significantly reduced within a week of treatment with ONC206 in mice bearing intracranial SF8628 tumors at 3T. Of note, there was no change in MRI-detectable tumor volume at day 7 (Supplementary Fig. S6F). Importantly, reduced lactate at day 7 reflected the ONC206-induced extension in survival at a later timepoint (62.5 days for ONC206-treated mice vs. 49 days for vehicle-treated controls, p<0.05; Fig. 7G).

We then questioned whether DMI provides an early readout of treatment response that reflects molecular alterations at the tissue level. To address this question, rats were intracranially implanted with BT245 cells, and once tumors were MR-visible, this time point was considered day 0, and 2D CSI data was acquired after administration of [6,6-^2^H]-glucose on a 9.4T scanner with a spatial resolution of 70.3 μL. Rats were then treated with ONC206 and 2D CSI data was again acquired on day 7±1. Rats were euthanized after the DMI scan, and tumor tissue was isolated for *ex vivo* analysis of pharmacodynamic biomarkers of treatment response (see the experimental schematic in Fig. 7H). Visualization of the data in the form of metabolic heatmaps generated by overlaying the SNR of lactate over the corresponding T2-weighted MRI showed a significant reduction in [3,3’-^2^H]-lactate localized to the tumor at day 7 relative to day 0 in BT245 tumor-bearing rats *in vivo* (Fig. 7I). Quantification of the data across multiple rats confirmed the statistically significant drop in [3,3’-^2^H]-lactate production from [6,6-^2^H]-glucose at day 7 relative to day 0 (Fig. 7J). To assess whether [3,3’-^2^H]-lactate was a pharmacodynamic biomarker of response to therapy, we examined tumor tissue resected from the ONC206-treated rats after the DMI scan. As controls, we examined tumor tissue resected from a separate cohort of vehicle-treated BT245 tumor-bearing rats. Since the imipridones act by hyperactivating the mitochondrial protease ClpP and inducing the degradation of mitochondrial proteins such as ClpX (6–12), we used flow cytometry to quantify ClpX expression in tumor tissue. As shown in Fig. 7K and Supplementary Fig. S6G, ClpX expression was significantly reduced in ONC206-treated tumors relative to vehicle-treated controls. These results indicate that quantifying lactate production from [6,6-^2^H]-glucose provides an early readout of drug-target engagement in DMGs *in vivo*.

To further confirm these results, we examined whether DMI of [6,6-^2^H]-glucose metabolism provides an early readout of response to ONC206 at the clinically relevant field strength of 3T (see schematic in Fig. 7L). Briefly, mice were intracranially implanted with BT245 cells, and once tumors were observed on T2-weighted MRI, mice were treated with either vehicle or ONC206. On day 7, 2D CSI data was acquired after administration of [6,6-^2^H]-glucose on a 3T scanner with a spatial resolution of 113.5 μL. Mice were then longitudinally treated with vehicle or ONC206 until they had to be euthanized according to animal care guidelines. As shown in the metabolic heatmaps in Fig. 7M and the quantification in Fig. 7N, [3,3’-^2^H]-lactate production from [6,6-^2^H]-glucose was significantly reduced in the tumor but not contralateral normal brain in ONC206-treated mice relative to controls at day 7. Of note, there was no significant difference in MRI-detectable tumor volume at this point (Supplementary Fig. S6H). Importantly, this reduction in [3,3’-^2^H]-lactate production preceded the significant extension in survival observed after ∼2 months (51 days for vehicle-treated controls vs. 62 days for ONC206-treated mice, p<0.005; Fig. 7O).

Finally, we examined whether DMI can assess early response to standard-of-care radiation *in vivo* (Fig. 7P). To this end, we implanted mice with DIPG-13 cells, and once tumors were MR-visible, mice were divided into control (untreated) or radiation (IR; 2 Gy/day for 3 days) groups. On day 3, 2D CSI data was acquired on a 3T scanner with a spatial resolution of 113.5 μL after administration of [6,6-^2^H]-glucose. Mice were euthanized after the DMI scan, and tumor tissue was resected for *ex vivo* analysis of tissue biomarkers of response to radiation (see Fig. 7P). As shown in Fig. 7Q-7R, [3,3’-^2^H]-lactate production from [6,6-^2^H]-glucose was significantly reduced in the tumor but not contralateral normal brain in mice subjected to radiation relative to controls at day 3. Concomitantly, flow cytometric analysis of *ex vivo* tumor tissue showed that phosphorylation of H2AX, which is a well-known biomarker of radiation response (33), was significantly higher in radiation-treated tumor tissue relative to vehicle controls at day 3 (Fig. 7S and Supplementary Fig. S6I). We also confirmed that caspase activity was significantly higher in radiation-treated tumor tissue relative to vehicle controls at day 3 (Supplementary Fig. S6J). Collectively, our studies indicate that visualizing lactate production from [6,6-^2^H]-glucose provides an early readout of response to standard and targeted therapy that precedes extended survival and reflects pharmacodynamic alterations at the tissue level in preclinical DMG models *in vivo* at the clinically relevant field strength of 3T.

## DISCUSSION

DMGs are universally fatal primary brain tumors in children that are driven by the H3K27M mutation (1–3). These diffusely infiltrative tumors occur in midline locations of the brain that prevent safe surgical resection. Therapeutic options are limited with no effective chemotherapies and standard-of-care radiation being largely palliative (6). Unraveling the unique biological consequences of the H3K27M mutation has the potential to identify novel therapeutic opportunities and disease biomarkers for DMGs. In this study, we demonstrate that the H3K27M mutation upregulates the glycolytic enzyme PGK1 and drives lactate production which, in turn, facilitates NTP biosynthesis, DNA replication, and tumor proliferation by lactylating NME1 (see schematic in Fig. 5Q). Importantly, we leverage this information to demonstrate that lactate production from [6,6-^2^H]-glucose metabolism is a quantitative imaging biomarker of the H3K27M mutation and early response to therapy in preclinical DMG models *in vivo*.

Oncogenic events reprogram metabolism to preferentially shunt glucose via glycolysis to lactate rather than via the TCA cycle, even under conditions of normal oxygen tension (15,16). This phenomenon, also known as aerobic glycolysis or the Warburg effect, is considered a metabolic hallmark of cancer (15,16). Although less energy efficient, the Warburg effect is considered to promote tumor growth via regeneration of NAD+ and maintenance of a pool of metabolic intermediates for biosynthesis (16). In addition, the original view of lactate as a waste product of glycolysis has given way to a nuanced understanding of lactate as an “oncometabolite” that facilitates tumor growth by acidifying the tumor microenvironment and serving as a fuel under nutrient-poor conditions (38). Here, using isogenic and patient-derived DMG models and patient biopsies, we show that the H3K27M mutation, which is the disease-defining oncogenic event in DMGs, drives lactate production in DMGs. Mechanistically, we show that the H3K27M mutation upregulates the rate-limiting glycolytic enzyme PGK1 via reduced H3K27me3 at the PGK1 promoter. Importantly, we show that lactate directly facilitates tumor proliferation by driving NTP synthesis. Rapidly dividing tumor cells need a steady supply of purine and pyrimidine nucleotides for progression through the S phase of the cell cycle (39). As a result, *de novo* nucleotide synthesis is stringently regulated in cancer cells (39). Our results indicate that lactate is essential for the synthesis of NTPs in DMG cells. Depleting lactate via orthogonal approaches i.e., silencing PGK1 and glucose starvation, selectively abrogates NTP synthesis, blocks cell cycle progression, and induces replication stress. Although it is possible that lactate also promotes DMG growth via the other mechanisms outlined above such as NAD+ regeneration and serving as a metabolic fuel, nevertheless, our studies demonstrate a role for lactate in promoting DMG proliferation via NTP synthesis.

We demonstrate, to the best of our knowledge for the first time, that lactate promotes NTP synthesis via lysine lactylation of the NDPK enzyme NME1. Lactylation is a post-translational modification that was recently discovered for its role in modulating gene expression in macrophages via lactylation of histones (19,20). Recent studies have extended the role of histone lactylation to tumor cells, including gliomas (40). Emerging studies have also identified lactylation of lysine residues on proteins, including metabolic enzymes, as a post-translational modification (19,20). However, the role of lactylation in DMG biology has not been explored. In this study, we demonstrate a hitherto unknown role for lactate in upregulating NDPK activity via lactylation of NME1. Since our studies indicated that lactate depletion specifically blocked the synthesis of NTPs (GTP, UTP, and CTP) but not NDPs (GDP, UDP, and CDP), we examined NDPK activity, which is responsible for the conversion of NDPs to NTPs (34). Using multiple methods, including co-immunoprecipitation and PLA, we show that lactate post-translationally upregulates NME1 activity via lysine lactylation in patient-derived DMG cells. Systematic examination of NME1 mutants in which each lysine residue is mutated to arginine (which cannot be lactylated) indicates that lactylation of NME1 at K49 is essential for NME1 activity, NTP synthesis, and tumor proliferation in patient-derived DMG cells. Importantly, examination of patient biopsies indicates that NME1 lactylation, NDPK activity, and NTP abundance are significantly higher in H3K27M-mutated tumors relative to H3 wild-type and normal brain, underscoring the clinical relevance of our results. Although further studies are needed to identify the lactyltransferase that mediates NME1 lactylation, nevertheless, our studies identify a unique H3K27M-lactate-NME1 cascade that is essential for tumor proliferation in DMGs.

NME1 has long been studied for its role as a metastasis suppressor gene because NME1 transcript levels are reduced in metastatic melanoma cells relative to primary melanoma (41). A negative correlation between NME1 mRNA and/or protein expression and metastatic potential has been identified in many cancers, including breast, head and neck, liver, and ovarian cancers (41). In contrast, a positive correlation between NME1 mRNA and/or protein and metastatic potential has been identified in lymphoma and neuroblastoma (42). However, whether NME1 promotes proliferation and the mechanisms by which it is regulated in primary brain tumors, including DMGs, is unclear. Our studies definitively indicate that NME1 activity is critical for the maintenance of NTP levels in DMGs. In this context, it is important to note that our studies indicate that NME1 activity, but not mRNA or protein expression, is upregulated via lactylation in DMG cells, thereby pointing to the importance of focusing on enzyme activity as opposed to mRNA or protein expression. Further studies are needed to determine whether targeting NME1 directly or blocking NME1 lactylation represents a therapeutic opportunity for DMGs.

In this study, we leverage the mechanistic association between the H3K27M mutation, lactate, NME1, and tumor proliferation to demonstrate that DMI of [6,6-^2^H]-glucose metabolism facilitates quantification of tumor burden and early response to therapy in DMGs. The Warburg effect has long been exploited for tumor imaging as exemplified by the widespread use of 2-[^18^F]-fluoro-2-deoxy-D-glucose positron emission tomography (FDG-PET) for response assessment in cancer (21). FDG-PET measures the high glucose uptake that is characteristic of the Warburg effect but does not provide a readout of downstream metabolism to lactate (21). Furthermore, FDG-PET suffers from poor contrast in brain tumors due to the high glucose uptake in the normal brain (21). We demonstrate that silencing the H3K27M mutation abrogates lactate production from [6,6-^2^H]-glucose in patient-derived DMG cells, thereby indicating that [6,6-^2^H]-glucose provides a direct readout of the H3K27M mutation. ^2^H-lactate production is significantly reduced following treatment with standard-of-care radiation or emerging experimental therapies such as ONC201, ONC206, and paxalisib in DMG cells. Importantly, using both mice and rats bearing intracranial tumor xenografts from multiple patient-derived DMG models, we show that spatially mapping lactate production from [6,6-^2^H]-glucose demarcates tumor from surrounding normal brain with excellent tumor-to-background contrast *in vivo.* Furthermore, DMI of glycolytically active tumor tissue provides a readout of response to standard-of-care radiation and targeted therapy *in vivo* within a few days of treatment, in the absence of MRI-detectable volumetric alterations. Of note, lactate production from [6,6-^2^H]-glucose is a quantitative pharmacodynamic imaging biomarker that reflects treatment-induced molecular alterations at an early time point that precedes extended survival *in vivo*. In essence, our studies establish, for the first time, the utility of DMI for imaging tumor burden and obtaining an early readout of response to therapy in preclinical DMG models.

Regarding clinical translation, our DMI studies were performed at both 9.4T and the widely available clinical magnetic field strength of 3T. Although a higher magnetic field strength is undeniably beneficial for DMI (43), our findings highlight the feasibility of robustly quantifying tumor burden and response to therapy at 3T, which sets the stage for widespread clinical dissemination. Of note, although [6,6-^2^H]-glucose was intravenously administered in our preclinical models, it is given orally to human volunteers and adult glioblastoma patients in clinical trials, which further facilitates clinical translation (23,27). Given the known difficulties of reliably assessing response to therapy using MRI in DMG patients (13,14), our studies provide a strong rationale for assessing whether visualizing metabolically active tumor tissue using DMI provides an early readout of response to therapy in DMG patients.

In summary, we demonstrate that the oncogenic H3K27M mutation fundamentally rewires glycolysis to drive proliferation in DMGs. H3K27M epigenetically upregulates the rate-limiting glycolytic enzyme PGK1 and facilitates lactate production. Lactate, in turn, activates NME1 by lysine lactylation, thereby driving NTP biosynthesis and tumor proliferation. Importantly, imaging glycolytic lactate production using DMI provides a readout of tumor proliferation and early response to standard-of-care and experimental therapies in clinically relevant patient-derived DMG models. Clinical translation of our studies has the potential to enable precision imaging of tumor burden and treatment response for DMG patients.

## MATERIALS AND METHODS

### Cell culture

BT245 (male; RRID: CVCL_IP13) and DIPG-13 (female; RRID: CVCL_IT41) parental (K27M) and H3.3K27M KO cells were a kind gift from Dr. Nada Jabado (28). SF8628 (female; RRID: CVCL_IT46) cells were purchased from Sigma Aldrich. SU-DIPG-6 (here referred to as DIPG-6; female; RRID: CVCL_IT40) cells were a kind gift from Dr. Michelle Monje. QCTB-R059 cells were a kind gift from Dr. Simon Robinson. 24-B7 and 24-D3 cells were a kind gift from Dr. Phoenix (30). All cell lines were maintained as neurospheres in Neurobasal-A medium (Gibco) containing 25 mM D-glucose, 2 mM L-glutamine, 2% B27 (Gibco), 1% N2 (Gibco), 100 U/mL penicillin and streptomycin, and 20 ng/mL each of human EGF, FGF, PDGF-AA, and PDGF-BB. To induce glucose starvation, cells were cultured in serum-free medium as described above for 72 h, except that the glucose concentration was reduced to 2.5 mM. For rescue, the medium was supplemented with 1 mM sodium lactate, 1 mM sodium citrate, or 500 μM dimethyl α-ketoglutarate for 72 h. Cell lines were routinely tested for mycoplasma contamination, authenticated by short tandem repeat fingerprinting (Cell Line Genetics), and assayed within 6 months of authentication.

### Gene silencing and overexpression

BT245 and DIPG-13 KO cells were generated by introducing frameshift deletions in the H3.3K27M mutant allele by CRISPR-Cas9 editing by the Jabado laboratory as described previously (28). To silence H3.3K27M in the DIPG-6, SF8628, and QCTB-R059 models, we used an ASO-based approach that has been described earlier (5). Specifically, we treated DMG cells with AUM*silence^TM^* ASOs (AUM Biotech) that were designed to target the mutant H3.3K27M allele. A 21-nucleotide ASO sequence (5’-AGGCGCACTCATGCGAGCGGC-3’) that is complementary to the mutant *H3F3A* allele and spans the mutation site in exon 2 was used (5). A scrambled sequence (5’-CCTTCCCTGAAGGTTCCTCC-3’) was used as control. These ASOs harbor a phosphorothioate backbone and next-generation sugar nucleotide modifications that enhance stability and enable efficient cellular uptake without the need for transfection reagents (44). The ASOs were added to the cell culture medium at a concentration of 5 μM. Loss of the mutant H3.3K27M protein was confirmed at 72 h by western blotting. To silence PGK1 in DMG cells, we used Edit-R All-in-one lentiviral particles expressing sgRNA targeting PGK1 and the Cas9 nuclease under the control of the hEF1a promoter and harboring a puromycin resistance marker (sgPGK1-1: Dharmacon, VSGH12609-256212517 and sgPGK1-2: Dharmacon, VSGH12609-256212522). Lentiviral particles expressing a non-targeting control sequence (sgNT: Dharmacon, VSGC11964) were used as controls. DMG cells were transduced with the lentivirus in serum-free medium using polybrene and selected using puromycin. PGK1 knockdown was verified using quantitative PCR (QPCR) and immunoblotting. To generate NME1 in which each lysine is mutated to arginine, we performed site-directed mutagenesis on the FLAG-NM23-H1 plasmid (Addgene, 25000) using the Phusion Site-Directed Mutagenesis Kit (ThermoFisher Scientific) according to the manufacturer’s instructions. Expression was verified by western blotting for the FLAG tag.

### QPCR

Gene expression was measured by QPCR on a QuantStudio 5 system (ThermoFisher Scientific), and data was normalized to β-actin. Total RNA was extracted using the TRIzol reagent and converted to cDNA using the High-Capacity cDNA Reverse Transcription Kit (ThermoFisher Scientific). QPCR was performed using the TaqMan Fast Advanced Master Mix (ThermoFisher Scientific) and the following primers: *PGK1* (forward: CCGCTTTCATGTGGAGGAAGAAG; reverse: CTCTGTGAGCAGTGCCAAAAGC); *SLC16A3* (forward: CCACAAGTTCTCCAGTGCCATTG; reverse: CGCCAGGATGAACACGTACATG); *NME1* (forward: ACCATCCGTGGAGACTTCTGCA; reverse: ACCAGTTCCTCAGGGTGAAACC); and β-actin (forward primer: AGAGCTACGAGCTGCCTGAC; reverse primer: AGCACTGTGTTGGCGTACAG).

### ChIP-QPCR

Chromatin was immunoprecipitated using an antibody specific to the H3K27me3 moiety (Active Motif, #39155), H3K18la moiety (PTMBio, PTM-1406RM), or rabbit IgG (Cell Signaling, #2729) and the High-Sensitivity ChIP Kit (Abcam, ab185913) according to manufacturer’s instructions. QPCR was then performed for *PGK1* or *NME1* using the primers described above. Data was expressed as fold enrichment relative to IgG control.

### Immunoprecipitation

∼1 x 10^7^ cells or ∼10 mg of tissue was lysed in RIPA buffer (25mM Tris-HCl pH 7.6, 150 mM NaCl, 1% NP-40, 1% sodium deoxycholate, 0.1% SDS) containing 0.5 mM phenylmethyl sulphonyl fluoride, 150 nM aprotinin and 1 μM each of leupeptin and E64 protease inhibitor. Lysates were cleared by centrifugation at 14,000 rpm for 15 min at 4 °C. The supernatant was pre-cleared with protein G agarose beads (Santa Cruz Biotechnology), incubated with anti-NME1 antibody (ThermoFisher Scientific, UM800025CF) or isotype IgG1 control (ThermoFisher Scientific, MOPC-21) overnight at 4°C followed by incubation with protein G agarose beads at 4°C for 2 h. Beads were washed 5 times with phosphate-buffered saline (PBS), and bound proteins were eluted by boiling in SDS-PAGE sample buffer (95°C for 10 min). Immunoprecipitated proteins (IP) and original lysate (input) were examined by immunoblotting as described below.

### Immunoblotting

Lysates or immunoprecipitated proteins were prepared as described above. Total cellular protein (∼20 µg) was separated on a 4-20% polyacrylamide gel (Bio-Rad) by sodium dodecyl sulfate-polyacrylamide gel electrophoresis and transferred onto Immobilon-FL PVDF membrane (Millipore). Membranes were blocked overnight in 5% milk (Santa Cruz Biotechnology) in TBST (20 mM Tris-HCl, pH 7.5, 500 mM NaCl, 0.1% Tween 20) at 4°C. Membranes were then washed 3 times for 5 min each in TBST and incubated with primary antibodies diluted in TBST for 1 h at room temperature. Following 3 washes of 10 min each with TBST, HRP-conjugated secondary antibodies were added for 1 h in TBST at room temperature. Membranes were washed thrice in TBST for 10 min each and developed using a Chemisolo imager (Azure Biosciences) and an enhanced chemiluminescence substrate kit (ThermoFisher Scientific). Blots were probed for H3K27M (Cell Signaling, 74829), global H3K27me3 (Cell Signaling, 9733), global H3K27Ac (Cell Signaling, 8173), total H3.3 (ThermoFisher Scientific, MA5-24667), PGK1 (Abcam, ab199438), NME1 (ThermoFisher Scientific, UM800025CF), pan-lactyl lysine (PTMBio, PTM-1401RM), and FLAG (Cell Signaling, 2368). β-actin (Cell Signaling, 4970) and Hsp90 (Cell Signaling, #4877) were used as loading controls. Goat anti-rabbit IgG-HRP (Cell Signaling, 7074) and horse anti-mouse IgG-HRP (Cell Signaling, 7076) were used as secondary antibodies.

### EdU assay

Cells were seeded in 6-well plates at a density of 3×10^5^ cells/well and subjected to glucose starvation and metabolite rescue for 72 h as described above. To quantify the proportion of EdU+ cells in the S phase, 1×10^6^ cells were stained with 10 μM EdU for 30 minutes. Cells were washed once with stain buffer (BD Biosciences) and fixed using the Cytofix/Cytoperm solution (BD Biosciences) for 15 minutes in the dark at 4°C. Following fixation, cells were washed once with Perm/Wash buffer (BD Biosciences) and incubated with the Click-iT™ Plus reaction cocktail (Click-iT™ Plus EdU Alexa Fluor™ 647 Flow Cytometry Assay Kit, ThermoFisher Scientific, C10634) for 30 minutes in the dark at room temperature. Cells were then washed once with the Perm/Wash buffer and stained with 10 μM Vybrant DyeCycle Green stain (ThermoFisher Scientific, V35004) for 30 minutes in the dark at room temperature. Cells were analyzed by flow cytometry on a MACSQuant 10 analyzer (Miltenyi Biotec).

### Phosphoprotein flow cytometry

To quantify phosphoprotein expression, cells were fixed with Phosflow Fix Buffer I (BD Biosciences) for 10 minutes at 37°C, washed once with Phosflow Perm/Wash Buffer I (BD Biosciences), and resuspended in the Phosflow Perm/Wash Buffer I to a concentration of 1×10^6^ cells in 100 μL. Cells were stained with phospho-Chk1-S345-FITC (ThermoFisher Scientific, MA5-37021), phospho-RPA2-S33 (ThermoFisher Scientific, A300-246A), or phospho-H2AX-S139-eFluor™ 660 (ThermoFisher Scientific, 50-9865-42) at a dilution of 1:100 for 30 minutes at room temperature. For phospho-Chk1-S345 and phospho-H2AX-S139, cells were washed once with Phosflow Perm/Wash Buffer I, resuspended in stain buffer, and examined using the MACSQuant 10 flow cytometer. For phospho-RPA2-S33 analysis, following staining with the primary antibody, cells were washed 3 times with Phosflow Perm/Wash Buffer I, incubated with fluorochrome-conjugated secondary antibody (Goat anti-Rabbit IgG (H+L) Cross-Adsorbed Secondary Antibody, Alexa Fluor™ 700, ThermoFisher Scientific, A-21038) at a dilution of 1:100 for 30 minutes at room temperature. Cells were then washed once with Phosflow Perm/Wash Buffer I, resuspended in stain buffer, and examined using the MACSQuant 10 flow cytometer.

### NME1 flow cytometry

To quantify NME1 expression, 1×10^6^ cells were fixed using the Cytofix/Cytoperm solution for 15 minutes in the dark at 4°C. Following fixation, cells were washed and resuspended in 100 μL of stain buffer. Cells were stained with anti-NME1 primary antibody (ThermoFisher Scientific, UM800025CF) at a dilution of 1:100 and incubated at 37°C for 30 minutes. Cells were then washed 3 times with Perm/Wash buffer, resuspended in 100 μL of stain buffer, and stained with fluorochrome-conjugated secondary antibody (Donkey anti-Mouse IgG secondary antibody, Alexa Fluor™ Plus 647, ThermoFisher Scientific, A32787) at a dilution of 1:100 for 30 minutes at room temperature. Cells were then washed thrice with Perm/Wash Buffer, resuspended in stain buffer, and examined using the MACSQuant 10 flow cytometer.

### NDPK activity assay

NDPK activity was quantified as described previously using a coupled pyruvate kinase-lactate dehydrogenase assay with ATP as the phosphate donor and UDP as the phosphate acceptor (45). 1×10^7^ cells were lysed in 50 μL of ice-cold NDPK assay buffer (100 mM Tris-HCl, pH 7.5, 10 mM MgCl2, and 100 mM KCl) with protease inhibitor cocktail and centrifuged at 8,000 rpm at 4 °C for 10 minutes. Protein content was quantified and 10 μL of the lysate (containing ∼2 μg of protein) was added to a 990 μL reaction mixture containing 100 mM Tris-HCl, pH 7.5, 10 mM MgCl2, 100 mM KCl, 0.4 mM NADH, 6 mM ATP, 0.7 mM UDP, 4 mM phosphoenolpyruvate, and 10 U each of pyruvate kinase and lactate dehydrogenase. NADH absorbance was then measured at 340 nm using the Promega GloMax Discover spectrometer.

### Doubling time

Cells were seeded in a 96-well plate at 10,000 cells per well and treated as needed. After 4 days, cell number was measured using the MACSQuant 10 analyzer. The doubling time was calculated using the following formula: doubling time = [4 days × (ln2)] / [ln (day 4 cell count/day 0 cell count)].

### Ex vivo analysis of tumor tissue

For all studies, tumor, and normal brain tissue were dissociated into single cells using the tumor dissociation kit (Miltenyi Biotec) according to the manufacturer’s instructions. NDPK activity was quantified as described above. To quantify EdU incorporation, single cells were stained with 10 μM EdU for 30 minutes, fixed and permeabilized, and EdU incorporation was measured using the Click-iT™ Plus EdU Alexa Fluor™ 647 Flow Cytometry Assay Kit, ThermoFisher Scientific, C10634) according to the manufacturer’s instructions. For all other cytometric assays, cells were fixed and permeabilized by resuspension in the Cytofix/Cytoperm solution followed by incubation in the dark at 4°C for 20 min. Cells were washed once with the Perm/Wash buffer before staining with the relevant antibodies. To quantify ClpX expression cells were stained with the ClpX antibody (ThermoFisher Scientific, PA5-79052) at a dilution of 1:250 for 30 minutes in the dark at room temperature. Cells were then washed 3 times with Perm/Wash buffer, resuspended in 100 μL of BD^TM^ Stain buffer, and stained with fluorochrome-conjugated secondary antibody (Goat anti-Rabbit IgG secondary antibody, Alexa Fluor™ 750, ThermoFisher Scientific, A-21039) at a dilution of 1:500 for 30 minutes at room temperature. Cells were washed 3 times with Perm/Wash Buffer resuspended in stain buffer and analyzed using the MACSQuant 10 flow cytometer. To measure phospho-H2AX, cells were stained for phospho-H2AX as described in the phosphoprotein flow cytometry section above.

### PLA

The Duolink flowPLA Mouse/Rabbit Starter Kit Far Red (Sigma-Aldrich, DUO94104) was used according to the manufacturer’s instructions (35–37). Briefly, cells were treated as needed and fixed and permeabilized by resuspension in the Cytofix/Cytoperm solution in the dark at 4°C for 20 min. Cells were washed once with Duolink wash buffer and resuspended in Duolink blocking solution at a concentration of 1×10^6^ cells in 100 μL. Following incubation at 37°C for 1 h, cells were pelleted and resuspended in 100 μL of Duolink antibody diluent containing the mouse anti-NME1 antibody (1:100) and the rabbit anti-pan-lactyl lysine antibody (1:100). For the negative isotype IgG controls, cells were resuspended in 100 μL of Duolink antibody diluent containing the mouse IgG1 isotype control (1:100; ThermoFisher Scientific, MA1-10407) and the rabbit IgG isotype control (1:100, PTMBio, PTM-5073). We also examined cells resuspended in each primary antibody alone as added controls. Following incubation at 37°C for 1 h, the cells were washed twice with Duolink wash buffer and resuspended in 100 μL of the Duolink antibody diluent containing the anti-mouse MINUS PLA probe and the anti-rabbit PLUS PLA probe at a 1:5 dilution. After incubation at 37°C for 1 h, the cells were washed twice with Duolink wash buffer and resuspended in 50 μL of ligation buffer (containing two bridging DNA oligonucleotides and a ligase at a 1:40 dilution). The DNA oligonucleotides hybridize with the two PLA probes, allowing the ligase to form a rolling circle of DNA when the probes are in close proximity (for example, when the primary antibodies are bound to a protein and a post-translational modification on the protein). Following incubation at 37°C for 30 minutes, the cells were washed twice in Duolink wash buffer and incubated with 50 μL of amplification solution (containing nucleotides, fluorescently labeled, complementary oligonucleotides, and DNA polymerase at a 1:80 dilution) for 100 minutes at 37°C. The DNA polymerase amplifies the circular DNA probe via rolling circle amplification, and the fluorescently labeled oligonucleotides hybridize to a specific sequence in the amplified DNA to yield an amplified fluorescence signal that is detectable by flow cytometry (35–37).. The cells were then washed twice in Duolink wash buffer, resuspended in stain buffer, and analyzed on a MACSQuant 10 flow cytometer with excitation at 644 nm and emission at 669 nm.

### Viability

The RealTime-Glo Assay (Promega) was used to measure live-cell viability according to the manufacturer’s instructions. The assay uses a viability NanoLuc substrate that can only be reduced by live cells. The reduced substrate diffuses into the medium and produces bioluminescence via a reaction with NanoLuc Luciferase in the media. 10,000 cells per well were seeded in a 96-well plate. After 72 h, the NanoLuc substrate and luciferase were added, and bioluminescence was measured using a Promega GloMax Discover instrument.

### Patient biopsies

Patient biopsies were obtained from the UCSF Brain Tumor Center Biorepository in compliance with the written informed consent policy. Biopsy use was approved by the Committee on Human Research at UCSF and research was approved by the Institutional Review Board at UCSF according to ethical guidelines established by the U.S. Common Rule.

### Stable isotope tracing in cells

3×10^5^ cells were seeded in 6-well plates and incubated in cell culture media in which glucose was replaced with 25 mM [U-^13^C]-glucose for 72 h. For H3.3K27M silencing experiments, cells were treated with the H3.3K27M ASO or scrambled ASO for 72 h and concomitantly incubated in [U-^13^C]-glucose-containing media. When needed, the medium was supplemented with 1 mM sodium lactate, 1 mM sodium citrate, or 500 μM dimethyl α-ketoglutarate for 72 h. Cells were washed with ice-cold ammonium acetate buffer (150 mM, pH 7.3). 1 ml of pre-cooled methanol/water (80:20 v/v) was added to each sample and incubated at -80 °C for 30 minutes. Samples were centrifuged at 14,000 rpm for 15 min at 4 °C to remove debris. The supernatant was lyophilized and reconstituted with 60 μL of pre-chilled acetonitrile/water (50:50, v/v), transferred into glass vials, and utilized for LC-MS as described below.

### LC-MS

LC-MS was performed using a Vanquish Ultra High-performance LC system coupled to an Orbitrap ID-X Tribrid mass spectrometer (ThermoFisher Scientific), equipped with a heated electrospray ionization (H-ESI) source capable of both positive and negative modes simultaneously (46). Before analysis, the MS instrument was calibrated using a calibration solution (FlexMix, ThermoFisher Fisher). Cell or tissue samples, along with blank controls, were placed in the autosampler. Individual samples were run alongside a pooled sample made from an equal mixture of all individual samples to ensure chromatographic consistency. Chromatographic separation of metabolites was achieved by hydrophilic interaction liquid chromatography (HILIC) using a Luna 3 NH2 column (150 mm x 2.1 mm, 3 μm, Phenomenex) in conjunction with a HILIC guard column (Phenomenex, 2.1 mm). The column temperature and flow rate were maintained at 27°C and 0.2 mL/min respectively. Mobile phases consisted of A (5 mM ammonium acetate, 48.5 mM ammonium hydroxide pH 9.9) and B (100% Acetonitrile). The following linear gradient was applied: 0.0-0.1 min: 85-80% B, 0.1-17.0 min: 80-5% B, 17.0-24.0 min: 5% B, 24.0-25.0 min: 5-85% B, 25.0-36.0 min: 85% B. The injection volume and auto sampler temperature were kept at 5 μL and 4°C respectively. High-resolution MS was acquired using a full scan method alternating between positive and negative polarities (spray voltages: +3800kV/-3100kV; sheath gas flow: 45 arbitrary units: auxiliary gas flow: 15 arbitrary units; sweep gas flow: 1 arbitrary unit; ion transfer tube temperature: 275°C; vaporizer temperature: 300°C). Three mass scan events were set for the duration of the 36-minute run time. The first was the negative polarity mass scan settings at full-scan-range; 70-975 m/z. Positive polarity mass scan settings were split to two scan events, full-scan-range; 70-360 m/z and 360-1500m/z, in that order. MS1 data were acquired at resolution of 60,000 with a standard automatic gain control and a maximum injection time of 100 ms. Data was acquired using Xcalibur software. Chromatograms were reviewed using FreeStyle (ThermoFisher Fisher) and a 5-ppm mass tolerance. Peak areas were quantified using TraceFinder from either positive or negative modes depending on previously run standards. Peak areas were corrected to blank samples and normalized to protein content and the total ion count of that sample. For ^13^C isotope tracing, % ^13^C labeling for each metabolite was calculated after correcting for natural abundance using Escher Trace (47).

### DMI of cell suspensions

For studies with K27M and KO cells, 2 x 10^6^ cells were seeded in T150 flasks and incubated in Neurobasal-A/B-27/N2-growth factor media in which glucose was replaced with 25 mM [6,6-^2^H]-glucose for 72 h. For radiation studies, cells were subjected to a single dose of 10 Gy radiation before incubation in [6,6-^2^H]-glucose-containing media for 72 h. For drug studies, cells were treated with vehicle (DMSO), 500 nM ONC206, 10 μM ONC201, 1 μM paxalisib, or the combination of 1 μM paxalisib and 5 μM ONC201 and concurrently incubated in [6,6-^2^H]-glucose-containing media for 72 h. These doses were chosen based on prior studies (9–11). At 72 h, cells were harvested, counted, and resuspended in regular medium. ^2^H-MR spectra were acquired on a Bruker 14.1T scanner using the zg2H sequence (TR = 1s, complex points=8192, NA=128, spectral width=1667 Hz, temporal resolution= 7 minutes 34 s). Peak integrals for lactate (1.3 ppm), glucose (3.75 ppm) and HDO (4.75 ppm) were calculated using Mnova v15 (MestreNova). The peak integral for lactate was normalized to that of glucose and HDO and to cell number (referred to as normalized lactate).

### Intracranial tumor implantation

All studies were performed in accordance with the National Institutes of Health Guide for the Care and Use of Laboratory Animals and study protocols were approved by the University of California, San Francisco Institutional Animal Care and Use Committee (IACUC). SF8628, BT245, or DIPG-13 cells (5×10^5^ cells in 3 μL) were intracranially injected into the cortex of SCID mice (female, 5-6 weeks old, Charles River Laboratories) as described previously (25,26). BT245, DIPG-6, or DIPG-13 cells were injected into the pons of SCID mice (5×10^5^ cells in 3 μL; female, 5-6 weeks old, Charles River Laboratories) or nude rats (5×10^5^ cells in 10 μL; male, 5-6 weeks old, Envigo) using the following coordinates from the lambda suture (x=0.8 mm, y=6.5 mm, z=1 mm).

### MRI

Tumor volume was determined by T2-weighted MRI using a small animal horizontal Bruker 3T or 9.4T scanner (25,26). For the 3T scanner, we used a ^1^H quadrature volume coil and a T2 rapid acquisition with relaxation enhancement (RARE) sequence (TE/TR = 64/3700 ms, FOV = 30 x 30 mm^2^, matrix = 256 x 256, slice thickness = 1.5 mm, NA = 5). Studies at 9.4T were performed using a ^1^H/^13^C linear/linear volume coil on a preclinical 9.4T MR scanner (Biospec, Bruker). Axial T2-weighted images were acquired using a spin-echo TurboRARE sequence (TE/TR = 8.25/3200ms, FOV = 30×30 mm^2^, 256×256, slice thickness=1.2 mm, NA=3).

### Optical imaging

For studies with DIPG-13 mice treated with radiation, tumor growth was monitored by bioluminescence on a Xenogen IVIS Spectrum 10 min after intraperitoneal injection of 150 mg/kg D-luciferin prepared according to the manufacturer’s directions (Gold Biotechnology, #LUCK-100). The total flux signal was used for quantification in all studies.

### DMI studies in vivo

DMI studies were performed on a small animal horizontal Bruker 3T or 9.4T scanner using a 16 mm ^2^H surface coil in combination with a quadrature ^1^H coil (40 mm inner diameter for the 3T scanner and 72 mm inner diameter for the 9.4T scanner). *Non-localized ^2^H-MR spectroscopy at 9.4T:* Once tumors were ∼35-50 mm^3^, mice bearing DIPG-6 tumors implanted in the pons were placed in the scanner and a pre-injection ^2^H-MR spectrum was acquired. Following intravenous injection of a bolus of 2 g/kg of [6,6’-^2^H]-glucose via a tail-vein catheter, non-localized ^2^H-MR spectra were acquired with a pulse-acquire sequence (TR=313.47ms, NA=1000, 256 points, flip angle=64, spectral width=819.67, temporal resolution = 5 min 13s). *Non-localized ^2^H-<u>MR spectroscopy at 3T:</u>* For studies with mice bearing BT245 xenografts implanted in the cortex, once tumors were ∼35-50 mm^3^, mice were placed in the scanner and non-localized ^2^H-MR spectra were acquired before and after intravenous administration of a bolus of 2 g/kg of [6,6’-^2^H]-glucose using a pulse-acquire sequence (TR=506.361 ms, NA=1000, complex points=256, flip angle=64, spectral width=512.8 Hz, temporal resolution=8 minutes 26 s). For studies with mice bearing SF8628 xenografts implanted in the cortex, when tumors were ∼35-50 mm^3^, mice were randomized and treated with vehicle (saline, twice daily) or ONC206 (25 mg/kg in saline, twice daily) for 5 days per week. After the acquisition of non-localized ^2^H-MR data on day 7 at 3T, mice were treated 5 days per week, and survival monitored longitudinally until they needed to be euthanized according to IACUC guidelines. *<u>2D CSI at 9.4T:</u>* 2D CSI was used to spatially map lactate distribution in mice bearing BT245 or DIPG-6 xenografts implanted in the pons and in rats bearing BT245 tumors implanted in the pons. Once tumors were ∼35-50 mm^3^ (mice) or ∼80 mm^3^ (rats), animals were placed in the Bruker 9.4T scanner, and a bolus of 2 g/kg of [6,6’-^2^H]-glucose was administered via a tail-vein catheter. 2D CSI data was acquired using the following parameters (TE/TR=1.104/591.824ms, FOV=30x30x5 mm^3^, 512 points, spectral width=2.5 kHz, NA=8, temporal resolution=8 minutes 30 s, nominal voxel size = 70.3 μL). To assess if DMI reports on early response to treatment with ONC206, 2D CSI data was acquired as described above from rats bearing pontine BT245 tumors when tumors were ∼80 mm^3^ (this time point was designated as day 0). Rats were then treated with ONC206 (25 mg/kg in saline, twice daily, 5 days/week) and 2D CSI data was again acquired at day 7±1 as described above. After the DMI scan, rats were euthanized, and tumor tissue was resected for *ex vivo* flow cytometric analysis of ClpX expression. *<u>2D CSI at 3T:</u>* These studies were performed on mice bearing intracranial BT245 or DIPG-13 tumors implanted in the cortex and in rats bearing intracranial DIPG-13 tumors implanted in the pons. 2D CSI data was acquired as described for the 9.4T with the following parameters (TE/TR = 1.04/265.89 ms, FOV = 30x30x8 mm^3^, complex points=128 points, spectral width=2.5 kHz, NA = 30, temporal resolution=8 minutes 30 s, nominal voxel size = 112.5 uL). To quantify [6,6’-^2^H]-glucose metabolism in response to radiation, mice bearing cortical DIPG-13 tumors were monitored by monitored by bioluminescence until tumors reached a radiance of ∼1×10^6^ p/s/cm^2^/sr. This time point was considered day 0 (D0) and mice were randomized into control (untreated) or radiation (2 Gy/day for 3 days) groups. Radiation was administered using a precision Rad320 irradiator equipped with an X-ray tube that can deliver X-rays up to 320 kVp/12.5 mA and an adjustable shelf for precise animal placement. After the final radiation dose on day 3, 2D CSI data was acquired after intraperitoneal administration of a bolus of 2 g/kg of [6,6’-^2^H]-glucose using the parameters described above. Mice were then euthanized, and tumor tissue was resected for flow cytometric quantification of phospho-H2AX expression. To examine [6,6’-^2^H]-glucose metabolism in response to treatment with ONC206, mice bearing intracranial cortical BT245 tumors of ∼35 mm^3^ were randomized and treated with vehicle (saline, twice daily) or ONC206 (25 mg/kg in saline, twice daily) for 5 days/week. On day 7, 2D CSI data was acquired at 3T using the parameters described above. Mice were then longitudinally treated for 5 days a week and monitored for survival until they needed to be euthanized according to IACUC guidelines. *<u>DMI data analysis:</u>* Nonlocalized ^2^H-MR spectra were analyzed using Mnova v15. The peak integral for lactate was normalized to the peak integral for glucose and pre-injection HDO (referred to as normalized lactate). 2D CSI data was analyzed using in-house Matlab codes as described earlier (25,26). For each voxel at every timepoint, peak integrals were calculated. To generate heatmaps of the SNR of ^2^H-lactate, raw data were interpolated from an 8x8 matrix to a 256 x 256 matrix and normalized to noise. The SNR of ^2^H-lactate was quantified from a volume of 10.99 mm^3^ volume placed over the tumor.

### Statistical analysis

All experiments were performed on a minimum of 3 biological replicates (n≥3), and the results were expressed as mean ± standard deviation. Statistical significance was assessed in GraphPad Prism 10 using a two-way ANOVA or two-tailed Welch’s t-test with p<0.05 considered significant. Analyses were corrected for multiple comparisons using Tukey’s method, wherever applicable. Survival was quantified using Kaplan-Meier analysis. * indicates statistical significance with p<0.05, ** indicates p<0.01, *** indicates p<0.001 and **** indicates p<0.0001.

### Data availability

The data in this manuscript is available from the corresponding author upon reasonable request.

## Acknowledgments

This study was funded by the following grants: ChadTough Defeat DIPG Foundation Game Changer grant P0567741 (PV, SM), Violet Foundation for Pediatric Brain Cancer (PV, SM), the National Institutes of Health grant R21CA289565 (PV), and the National Institutes of Health grant R01CA292674 (PV). The authors acknowledge support from the Preclinical NMR Imaging Core of the Department of Radiology and Biomedical Imaging at UCSF.

## Author’s Contributions

*Conceptualization, supervision, project administration, funding acquisition:* PV. *Development of methodology:* PV, GB. *Investigation:* PV, GB, CT, SU, AMG. *Formal Analysis:* PV, GB. *Software:* GB. *Resources:* CK, TP, SV, SPR, SM. *Writing – original draft:* PV. *Writing – review & editing:* PV.

## REFERENCES

1. Saratsis AM, Knowles T, Petrovic A, Nazarian J. H3K27M Mutant Glioma: Disease Definition and Biological Underpinnings. Neuro-oncology 2023 doi 10.1093/neuonc/noad164.

2. van den Bent M, Saratsis AM, Geurts M, Franceschi E. H3 K27M-altered glioma and diffuse intrinsic pontine glioma: Semi-systematic review of treatment landscape and future directions. Neuro-oncology 2024;26(Supplement_2):S110–s24 doi 10.1093/neuonc/noad220.

3. Haase S, Carney S, Varela ML, Mukherji D, Zhu Z, Li Y, et al. Epigenetic reprogramming in pediatric gliomas: from molecular mechanisms to therapeutic implications. Trends Cancer 2024;10(12):1147–60 doi 10.1016/j.trecan.2024.09.007.

4. Silveira AB, Kasper LH, Fan Y, Jin H, Wu G, Shaw TI, et al. H3.3 K27M depletion increases differentiation and extends latency of diffuse intrinsic pontine glioma growth in vivo. Acta Neuropathol 2019;137(4):637–55 doi 10.1007/s00401-019-01975-4.

5. Zhang Q, Yang L, Liu YH, Wilkinson JE, Krainer AR. Antisense oligonucleotide therapy for H3.3K27M diffuse midline glioma. Science translational medicine 2023;15(691):eadd8280 doi 10.1126/scitranslmed.add8280.

6. Jackson ER, Persson ML, Fish CJ, Findlay IJ, Mueller S, Nazarian J, et al. A review of current therapeutics targeting the mitochondrial protease ClpP in diffuse midline glioma, H3 K27-altered. Neuro-oncology 2024;26(Supplement_2):S136–s54 doi 10.1093/neuonc/noad144.

7. Arrillaga-Romany I, Miller JJ. Demonstrated efficacy and mechanisms of sensitivity of ONC201: H3K27M-mutant diffuse midline glioma in the spotlight. Neuro-oncology 2024:noae051 doi 10.1093/neuonc/noae051.

8. Bonner ER, Waszak SM, Grotzer MA, Mueller S, Nazarian J. Mechanisms of imipridones in targeting mitochondrial metabolism in cancer cells. Neuro-oncology 2021;23(4):542–56 doi 10.1093/neuonc/noaa283.

9. Venneti S, Kawakibi AR, Ji S, Waszak SM, Sweha SR, Mota M, et al. Clinical Efficacy of ONC201 in H3K27M-Mutant Diffuse Midline Gliomas Is Driven by Disruption of Integrated Metabolic and Epigenetic Pathways. Cancer discovery 2023:OF1–OF24 doi 10.1158/2159-8290.Cd-23-0131.

10. Jackson ER, Duchatel RJ, Staudt DE, Persson ML, Mannan A, Yadavilli S, et al. ONC201 in Combination with Paxalisib for the Treatment of H3K27-Altered Diffuse Midline Glioma. Cancer research 2023:Of1–of17 doi 10.1158/0008-5472.Can-23-0186.

11. Przystal JM, Cianciolo Cosentino C, Yadavilli S, Zhang J, Laternser S, Bonner ER, et al. Imipridones affect tumor bioenergetics and promote cell lineage differentiation in diffuse midline gliomas. Neuro-oncology 2022;24(9):1438–51 doi 10.1093/neuonc/noac041.

12. Arrillaga-Romany I, Gardner SL, Odia Y, Aguilera D, Allen JE, Batchelor T, et al. ONC201 (Dordaviprone) in Recurrent H3 K27M–Mutant Diffuse Midline Glioma. Journal of Clinical Oncology 2024;42(13):1542–52 doi 10.1200/JCO.23.01134.

13. Cooney TM, Cohen KJ, Guimaraes CV, Dhall G, Leach J, Massimino M, et al. Response assessment in diffuse intrinsic pontine glioma: recommendations from the Response Assessment in Pediatric Neuro-Oncology (RAPNO) working group. Lancet Oncol 2020;21(6):e330–e6 doi 10.1016/s1470-2045(20)30166-2.

14. Chiang J, Diaz AK, Makepeace L, Li X, Han Y, Li Y, et al. Clinical, imaging, and molecular analysis of pediatric pontine tumors lacking characteristic imaging features of DIPG. Acta Neuropathol Commun 2020;8(1):57 doi 10.1186/s40478-020-00930-9.

15. Venneti S, Thompson CB. Metabolic Reprogramming in Brain Tumors. Annu Rev Pathol 2017;12:515–45 doi 10.1146/annurev-pathol-012615-044329.

16. Lunt SY, Vander Heiden MG. Aerobic glycolysis: meeting the metabolic requirements of cell proliferation. Annual review of cell and developmental biology 2011;27:441–64 doi 10.1146/annurev-cellbio-092910-154237.

17. Pavlova NN, Zhu J, Thompson CB. The hallmarks of cancer metabolism: Still emerging. Cell metabolism 2022;34(3):355–77 doi 10.1016/j.cmet.2022.01.007.

18. Gantner BN, Palma FR, Pandkar MR, Sakiyama MJ, Arango D, DeNicola GM, et al. Metabolism and epigenetics: drivers of tumor cell plasticity and treatment outcomes. Trends Cancer 2024;10(11):992–1008 doi 10.1016/j.trecan.2024.08.005.

19. He Y, Song T, Ning J, Wang Z, Yin Z, Jiang P, et al. Lactylation in cancer: Mechanisms in tumour biology and therapeutic potentials. Clin Transl Med 2024;14(11):e70070 doi 10.1002/ctm2.70070.

20. Li H, Sun L, Gao P, Hu H. Lactylation in cancer: Current understanding and challenges. Cancer cell 2024;42(11):1803–7 doi 10.1016/j.ccell.2024.09.006.

21. Kim MM, Parolia A, Dunphy MP, Venneti S. Non-invasive metabolic imaging of brain tumours in the era of precision medicine. Nature Reviews Clinical Oncology 2016;13:725 doi 10.1038/nrclinonc.2016.108.

22. Ruiz-Rodado V, Brender JR, Cherukuri MK, Gilbert MR, Larion M. Magnetic resonance spectroscopy for the study of cns malignancies. Prog Nucl Magn Reson Spectrosc 2021;122:23–41 doi 10.1016/j.pnmrs.2020.11.001.

23. De Feyter HM, Behar KL, Corbin ZA, Fulbright RK, Brown PB, McIntyre S, et al. Deuterium metabolic imaging (DMI) for MRI-based 3D mapping of metabolism in vivo. Sci Adv 2018;4(8):eaat7314 doi 10.1126/sciadv.aat7314.

24. Lu M, Zhu XH, Zhang Y, Mateescu G, Chen W. Quantitative assessment of brain glucose metabolic rates using in vivo deuterium magnetic resonance spectroscopy. Journal of cerebral blood flow and metabolism: official journal of the International Society of Cerebral Blood Flow and Metabolism 2017;37(11):3518–30 doi 10.1177/0271678x17706444.

25. Taglang C, Batsios G, Mukherjee J, Tran M, Gillespie AM, Hong D, et al. Deuterium magnetic resonance spectroscopy enables noninvasive metabolic imaging of tumor burden and response to therapy in low-grade gliomas. Neuro-oncology 2022;24(7):1101–12 doi 10.1093/neuonc/noac022.

26. Batsios G, Taglang C, Tran M, Stevers N, Barger C, Gillespie AM, et al. Deuterium Metabolic Imaging Reports on TERT Expression and Early Response to Therapy in Cancer. Clinical cancer research: an official journal of the American Association for Cancer Research 2022;28(16):3526–36 doi 10.1158/1078-0432.Ccr-21-4418.

27. Kaggie JD, Khan AS, Matys T, Schulte RF, Locke MJ, Grimmer A, et al. Deuterium metabolic imaging and hyperpolarized (13)C-MRI of the normal human brain at clinical field strength reveals differential cerebral metabolism. Neuroimage 2022;257:119284 doi 10.1016/j.neuroimage.2022.119284.

28. Krug B, De Jay N, Harutyunyan AS, Deshmukh S, Marchione DM, Guilhamon P, et al. Pervasive H3K27 Acetylation Leads to ERV Expression and a Therapeutic Vulnerability in H3K27M Gliomas. Cancer cell 2019;35(5):782–97.e8 doi 10.1016/j.ccell.2019.04.004.

29. Bartman CR, Faubert B, Rabinowitz JD, DeBerardinis RJ. Metabolic pathway analysis using stable isotopes in patients with cancer. Nature Reviews Cancer 2023;23(12):863–78 doi 10.1038/s41568-023-00632-z.

30. du Chatinier A, Meel MH, Das AI, Metselaar DS, Waranecki P, Bugiani M, et al. Generation of immunocompetent syngeneic allograft mouse models for pediatric diffuse midline glioma. Neuro-Oncology Advances 2022;4(1):vdac079 doi 10.1093/noajnl/vdac079.

31. Diehl FF, Miettinen TP, Elbashir R, Nabel CS, Darnell AM, Do BT, et al. Nucleotide imbalance decouples cell growth from cell proliferation. Nature cell biology 2022;24(8):1252–64 doi 10.1038/s41556-022-00965-1.

32. Do BT, Hsu PP, Vermeulen SY, Wang Z, Hirz T, Abbott KL, et al. Nucleotide depletion promotes cell fate transitions by inducing DNA replication stress. Developmental Cell 2024;59(16):2203–21.e15 doi 10.1016/j.devcel.2024.05.010.

33. Saxena S, Zou L. Hallmarks of DNA replication stress. Mol Cell 2022;82(12):2298–314 doi 10.1016/j.molcel.2022.05.004.

34. Lacombe ML, Milon L, Munier A, Mehus JG, Lambeth DO. The human Nm23/nucleoside diphosphate kinases. Journal of bioenergetics and biomembranes 2000;32(3):247–58 doi 10.1023/a:1005584929050.

35. Fredriksson S, Gullberg M, Jarvius J, Olsson C, Pietras K, Gústafsdóttir SM, et al. Protein detection using proximity-dependent DNA ligation assays. Nature biotechnology 2002;20(5):473–7 doi 10.1038/nbt0502-473.

36. Sharanek A, Raco L, Soleimani VD, Jahani-Asl A. In situ detection of protein-protein interaction by proximity ligation assay in patient derived brain tumor stem cells. STAR Protocols 2022;3(3):101554 doi 10.1016/j.xpro.2022.101554.

37. Andersen SS, Hvid M, Pedersen FS, Deleuran B. Proximity ligation assay combined with flow cytometry is a powerful tool for the detection of cytokine receptor dimerization. Cytokine 2013;64(1):54–7 doi 10.1016/j.cyto.2013.04.026.

38. Lanzetti L. Oncometabolites at the crossroads of genetic, epigenetic and ecological alterations in cancer. Cell Death & Differentiation 2024;31(12):1582–94 doi 10.1038/s41418-024-01402-6.

39. Mullen NJ, Singh PK. Nucleotide metabolism: a pan-cancer metabolic dependency. Nature Reviews Cancer 2023;23(5):275–94 doi 10.1038/s41568-023-00557-7.

40. Wang S, Huang T, Wu Q, Yuan H, Wu X, Yuan F, et al. Lactate reprograms glioblastoma immunity through CBX3-regulated histone lactylation. The Journal of clinical investigation 2024;134(22) doi 10.1172/jci176851.

41. Hartsough MT, Steeg PS. Nm23/Nucleoside Diphosphate Kinase in Human Cancers. Journal of bioenergetics and biomembranes 2000;32(3):301–8 doi 10.1023/A:1005597231776.

42. Tan C-Y, Chang CL. NDPKA is not just a metastasis suppressor – be aware of its metastasis-promoting role in neuroblastoma. Laboratory Investigation 2018;98(2):219–27 doi 10.1038/labinvest.2017.105.

43. de Graaf RA, Hendriks AD, Klomp DWJ, Kumaragamage C, Welting D, Arteaga de Castro CS, et al. On the magnetic field dependence of deuterium metabolic imaging. NMR in biomedicine 2020;33(3):e4235 doi 10.1002/nbm.4235.

44. Rinaldi C, Wood MJA. Antisense oligonucleotides: the next frontier for treatment of neurological disorders. Nature Reviews Neurology 2018;14(1):9–21 doi 10.1038/nrneurol.2017.148.

45. Krishnan KS, Rikhy R, Rao S, Shivalkar M, Mosko M, Narayanan R, et al. Nucleoside Diphosphate Kinase, a Source of GTP, Is Required for Dynamin-Dependent Synaptic Vesicle Recycling. Neuron 2001;30(1):197–210 doi 10.1016/S0896-6273(01)00273-2.

46. Udutha S, Taglang C, Batsios G, Gillespie AM, Tran M, Ronen SM, et al. Telomerase reverse transcriptase induces targetable alterations in glutathione and nucleotide biosynthesis in glioblastomas. bioRxiv 2023 doi 10.1101/2023.11.14.566937.

47. Kumar A, Mitchener J, King ZA, Metallo CM. Escher-Trace: a web application for pathway-based visualization of stable isotope tracing data. BMC Bioinformatics 2020;21(1):297 doi 10.1186/s12859-020-03632-0.

