## Supplementary figures and legends for "Lactylation fuels nucleotide biosynthesis and facilitates deuterium metabolic imaging of tumor proliferation in H3K27M-mutant gliomas"

### Supplementary Figure 1

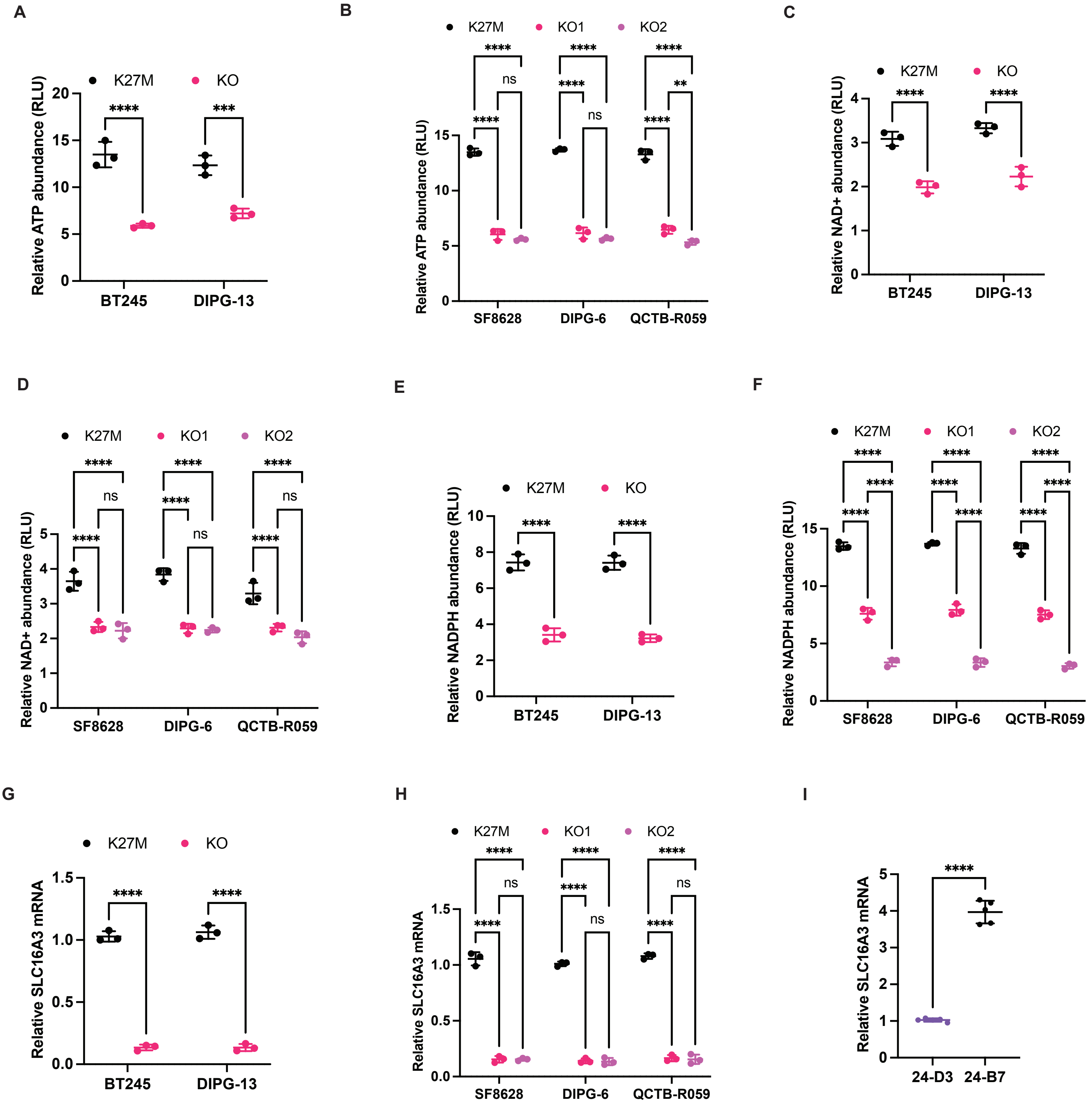

**Supplementary Figure 1. The H3K27M mutation rewires glucose metabolism in patient-derived DMG models. (A)** Quantification of ATP in K27M and KO cells for the BT245 and DIPG-13 models. **(B)** Quantification of ATP in K27M and KO cells for the SF8628, DIPG-6, and QCTB-R059 models. **(C)** Quantification of NAD+ in K27M and KO cells for the BT245 and DIPG-13 models. **(D)** Quantification of NAD+ in K27M and KO cells for the SF8628, DIPG-6, and QCTB-R059 models. **(E)** Quantification of NADPH in K27M and KO cells for the BT245 and DIPG-13 models. **(F)** Quantification of NADPH in K27M and KO cells for the SF8628, DIPG-6, and QCTB-R059 models. **(G)** Quantification of SLC16A3 expression in K27M and KO cells for the BT245 and DIPG-13 models. **(H)** Quantification of SLC16A3 mRNA in K27M and KO cells for the SF8628, DIPG-6, and QCTB-R059 models. **(I)** Quantification of SLC16A3 expression in isogenic 24-B7 and 24-D3 cells.

#### Supplementary Figure 2

A

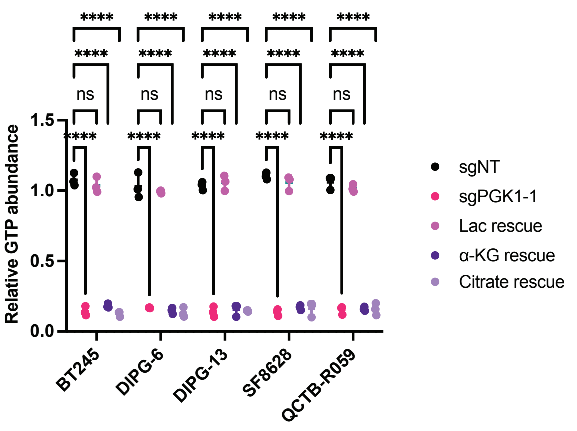


B

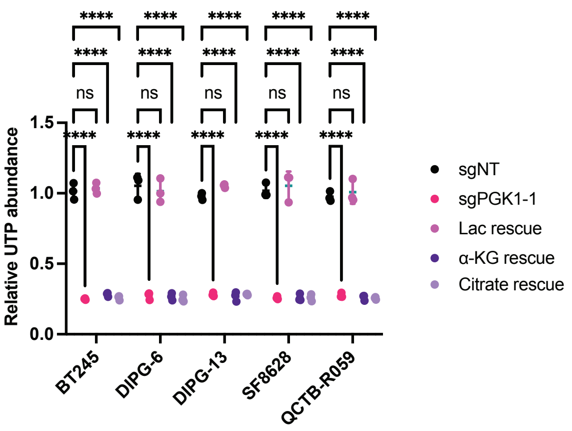


C

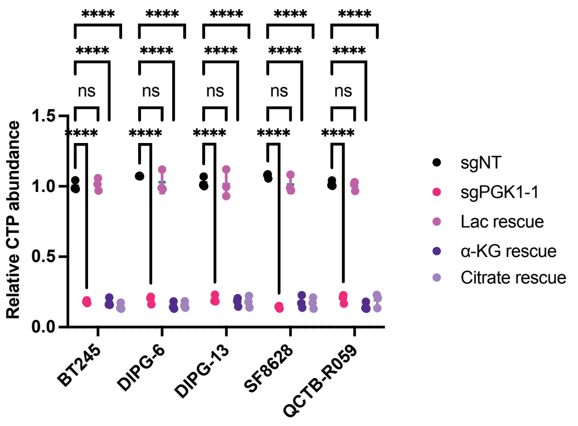

**Supplementary Figure 2. Lactate is essential for NTP synthesis in DMG cells.** Effect of supplementation with exogenous 1 mM sodium lactate (referred to as Lac rescue), 1 mM sodium citrate (Citrate rescue), or 500 μM dimethyl α-ketoglutarate (α-KG rescue) on the abundance of GTP **(A)**, UTP **(B)**, and CTP **(C)** in BT245, DIPG-13, SF8628, DIPG-6, and QCTB-R059 cells expressing sgRNA against PGK1 or a non-targeting control.

### Supplementary Figure 3

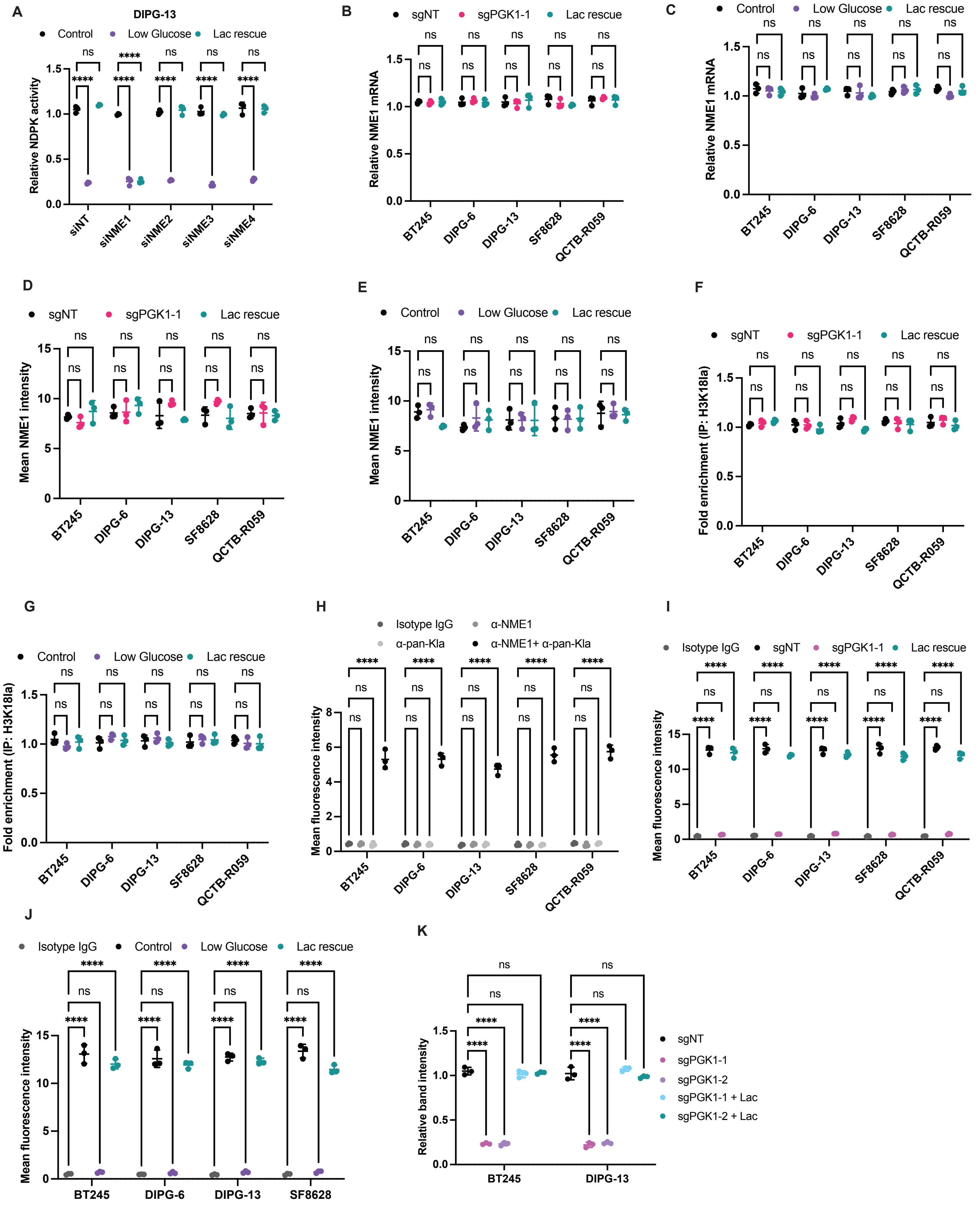

**Supplementary Figure 3. Lactate drives NTP synthesis by lactylating NME1. (A)** Effect of silencing NME1, NME2, NME3, or NME4 on the ability of lactate to rescue NDPK activity in DIPG-13 cells subjected to glucose starvation. **(B)** NME1 mRNA in sgNT, sgPGK1, and sgPGK1 cells treated with 1 mM sodium lactate for the BT245, DIPG-13, SF8628, DIPG-6, and QCTB-R059 models. **(C)** NME1 mRNA in BT245, DIPG-13, SF8628, DIPG-6, and QCTB-R059 cells subjected to glucose starvation and lactate rescue. **(D)** NME1 protein in sgNT, sgPGK1, and sgPGK1 cells treated with 1 mM sodium lactate for the BT245, DIPG-13, SF8628, DIPG-6, and QCTB-R059 models. **(E)** NME1 protein in BT245, DIPG-13, SF8628, DIPG-6, and QCTB-R059 cells subjected to glucose starvation and lactate rescue. **(F)** Fold enrichment of the histone H3K18 lactylation (H3K18la) mark at the NME1 promoter in sgNT, sgPGK1, and sgPGK1 cells treated with 1 mM sodium lactate for the BT245, DIPG-13, SF8628, DIPG-6, and QCTB-R059 models. **(G)** Fold enrichment of the histone H3K18 lactylation (H3K18la) mark at the NME1 promoter in BT245, DIPG-13, SF8628, DIPG-6, and QCTB-R059 cells subjected to glucose starvation and lactate rescue. **(H)** Quantification of NME1 lactylation by the PLA in BT245, DIPG-13, SF8628, DIPG-6, and QCTB-R059 cells. Cells were treated with a mixture of the mouse anti-NME1 and rabbit Pan-Kla antibodies (referred to as $\alpha$-NME1 + $\alpha$-Pan-Kla). Cells treated with the mixture of the corresponding isotype IgG antibodies (referred to as Isotype IgG mix) or with either primary antibody alone (referred to as $\alpha$-NME1 alone or $\alpha$-Pan-Kla alone) were used as negative controls. **(I)** Quantification of NME1 lactylation by the PLA in sgNT, sgPGK1, and sgPGK1 cells treated with 1 mM sodium lactate for the BT245, DIPG-13, SF8628, DIPG-6, and QCTB-R059 models. **(J)** Quantification of NME1 lactylation by the PLA in BT245, DIPG-13, SF8628, and DIPG-6 cells subjected to glucose starvation and lactate rescue. **(K)** Quantification of the effect of PGK1 silencing and lactate rescue on NME1 lactylation by immunoprecipitation and immunoblotting.

#### Supplementary Figure 4

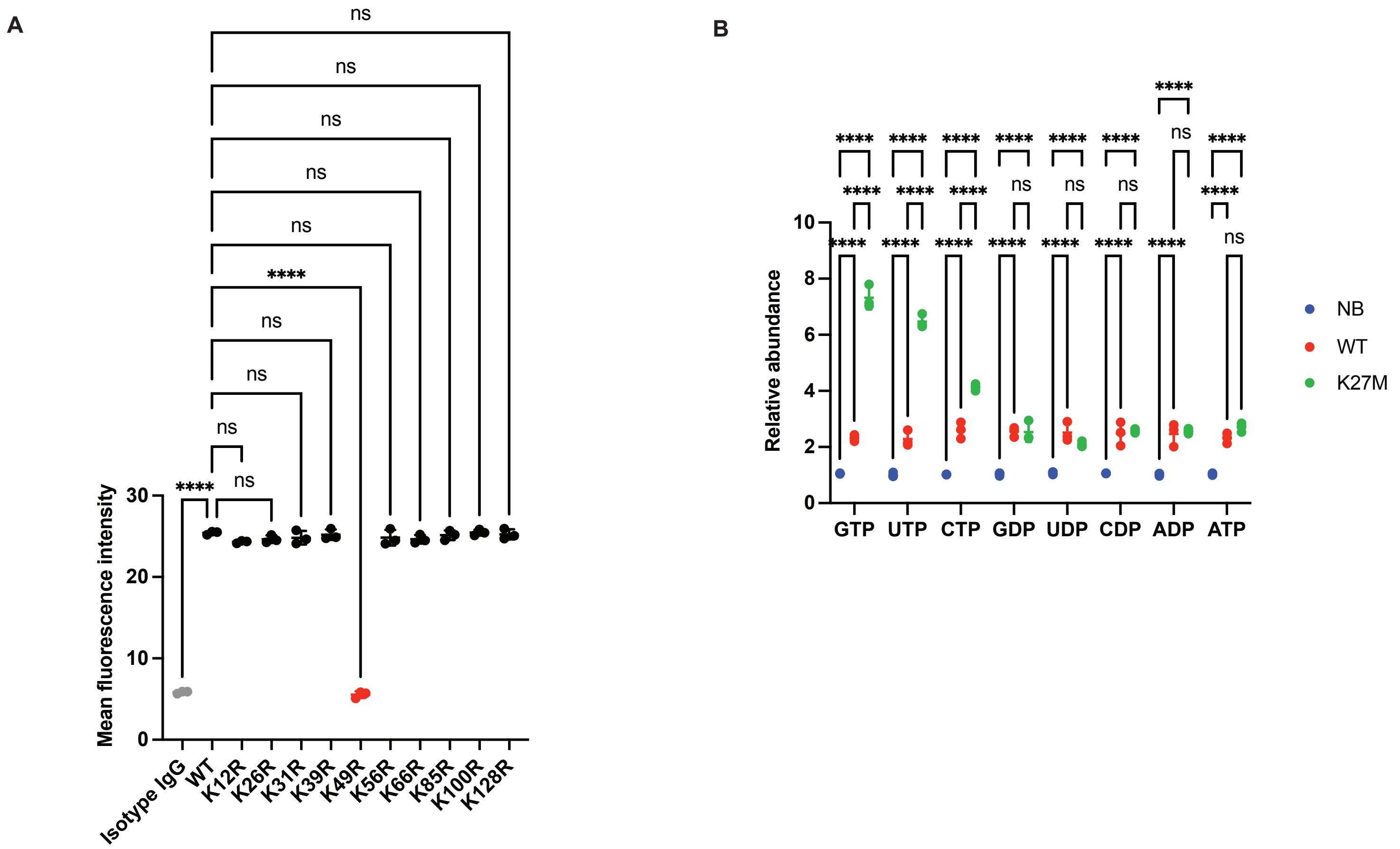

**Supplementary Figure 4. Lactate causes NME1 lactylation at K49 in DMG cells. (A)** Quantification of NME1 lactylation by the PLA in BT245 cells expressing wild-type (WT) NME1 or NME1 in which each lysine residue is mutated to arginine. Cells were treated with a mixture of the mouse anti-NME1 and rabbit Pan-Kla antibodies. Cells treated with the mixture of the corresponding isotype IgG were used as negative controls. **(B)** NTP and NDP abundance in biopsies from patients with H3K27M mutant (K27M), H3 wild-type (WT) tumors, or normal brain tissue (NB).

Supplementary Figure 5

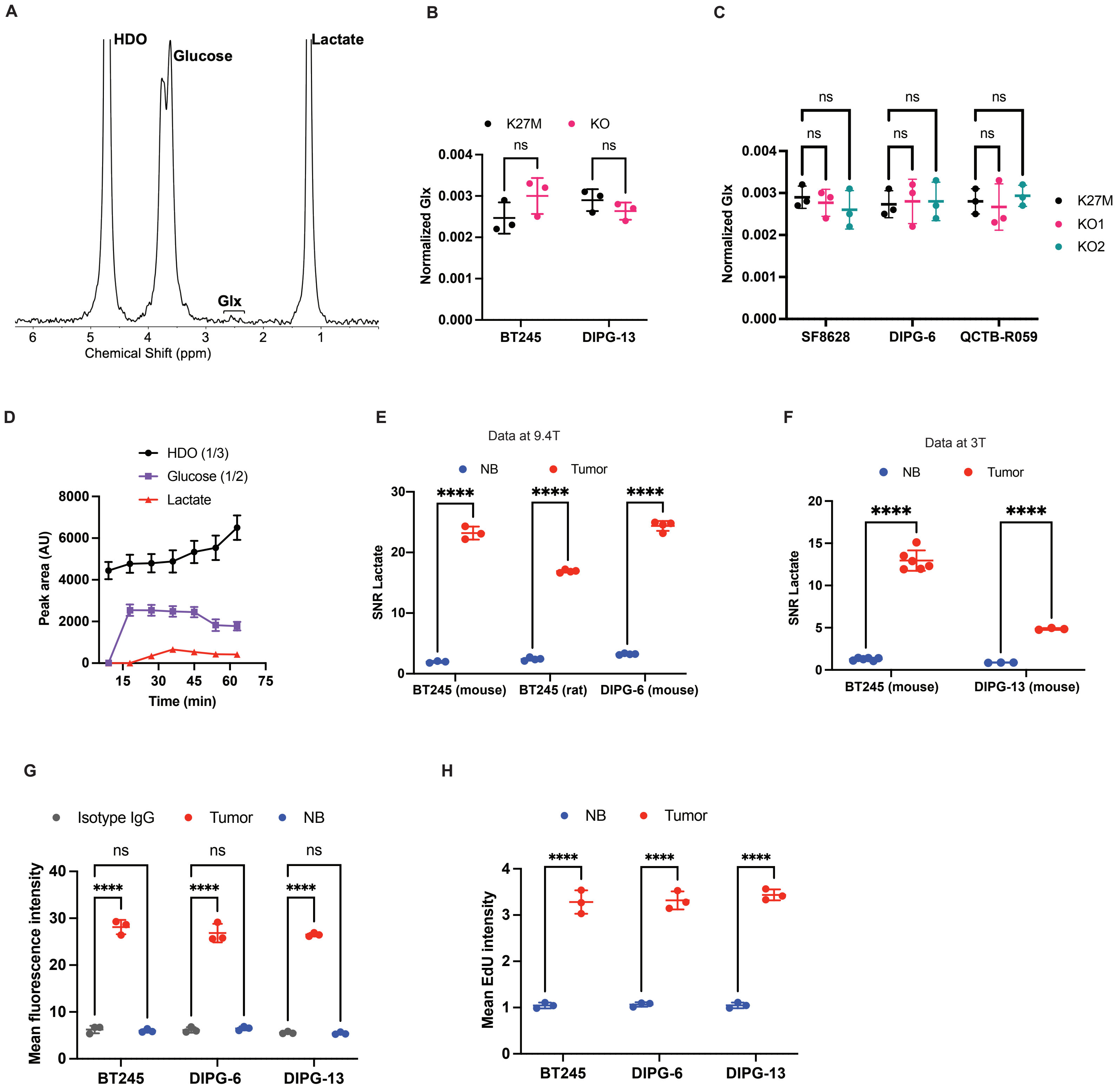

**Supplementary Figure 5. [6,6'-$^{2}$H]-glucose enables visualization of the metabolically active tumor lesion in preclinical DMG models. (A)** Representative $^{2}$H-MR spectra from BT245 K27M cells incubated in media containing 25 mM [6,6'-$^{2}$H]-glucose for 72 h. The peak for Glx at 2.4 ppm is highlighted. **(B)** Quantification of glx production from [6,6'-$^{2}$H]-glucose in K27M and KO cells for the BT245 and DIPG-13 models. **(C)** Quantification of glx production from [6,6'-$^{2}$H]-glucose in K27M and KO cells for the SF8628, DIPG-6, and QCTB-R059 models. **(D)** Quantification of the kinetics of HDO, glucose, and lactate following intravenous administration of [6,6'-$^{2}$H]-glucose into mice bearing intracranial BT245 tumors implanted in the cortex at 3T. **(E)** Quantification of the SNR of lactate in tumor and contralateral normal brain from 2D CSI data acquired at 9.4T in mice bearing intracranial cortical BT245 tumors, mice with intracranial DIPG-6 tumors in the pons, and rats bearing intracranial pontine BT245 tumors. **(F)** Quantification of the SNR of lactate in the tumor and contralateral normal brain from 2D CSI data acquired at 3T in mice bearing intracranial BT245 or DIPG-13 tumors implanted in the cortex. **(G)** Quantification of NME1 lactylation by the PLA **(G)** and EdU incorporation **(H)** in tumor and contralateral normal brain tissue resected from mice bearing BT245, DIPG-13, or DIPG-6 tumors.

#### Supplementary Figure 6

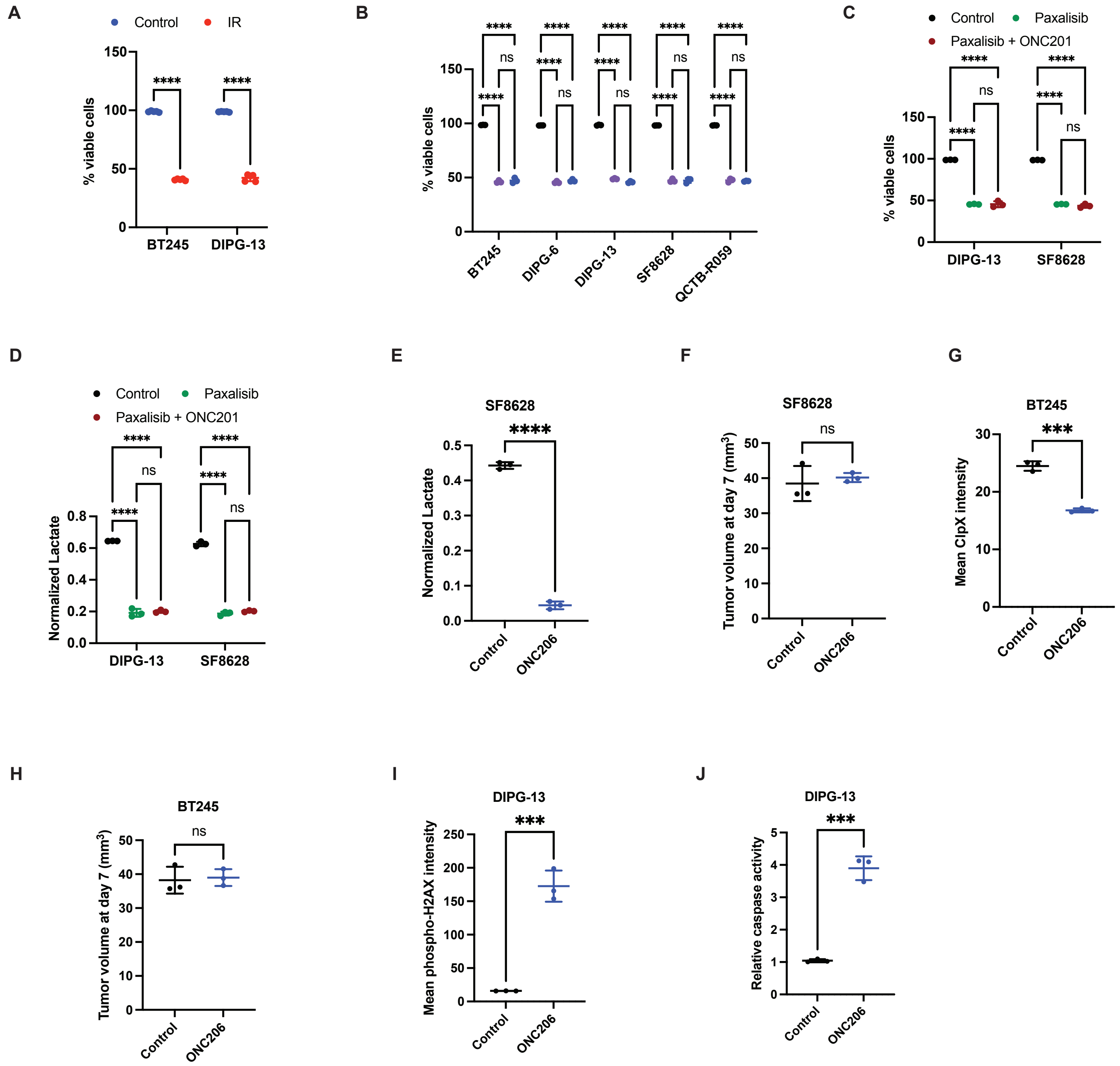

**Supplementary Figure 6. Lactate is a quantitative imaging biomarker of response to therapy in preclinical DMG models. (A)** Effect of 10 Gy IR on the viability of BT245 and DIPG-13 cells. **(B)** Effect of 10 μM ONC201 or 500 nM ONC206 on the viability of BT245, DIPG-13, SF8628, DIPG-6, and QCTB-R059 cells. **(C)** Effect of 1 μM paxalisib or the combination of 1 μM paxalisib and 5 μM ONC201 on the viability of SF8628 and DIPG-13 cells. **(D)** Effect of 1 μM paxalisib or the combination of 1 μM paxalisib and 5 μM ONC201 on [3,3'-$^{2}$H]-lactate production from [6,6'-$^{2}$H]-glucose in SF8628 and DIPG-13 cells. **(E)** Quantification of [3,3'-$^{2}$H]-lactate production from [6,6'-$^{2}$H]-glucose in mice bearing cortical SF8628 tumors treated with vehicle or ONC206 daily for 7 days. On day 7, $^{2}$H-MR data was acquired using a non-localized sequence following intravenous administration of [6,6'-$^{2}$H]-glucose into the mice at 3T. **(F)** Tumor volume at day 7 as measured by T2-weighted MRI in mice bearing cortical SF8628 tumors treated with vehicle or ONC206 for 7 days. **(G)** Quantification of ClpX expression in tumor tissue resected from BT245 tumor-bearing rats treated with ONC206 as described in Fig. 7H. Tumor tissue resected from a separate cohort of vehicle-treated rats was used as control. **(H)** Tumor volume at day 7 as measured by T2-weighted MRI in mice bearing intracranial BT245 tumors treated with vehicle or ONC206 for 7 days as described in Fig. 7L. Quantification of phospho-H2AX expression **(I)** and caspase activity **(J)** in tumor tissue resected from DIPG-13 control or IR-treated mice at day 3 (see Fig. 7P).
